# PP1 phosphatases control PAR-2 localization and polarity establishment in *C. elegans* embryos

**DOI:** 10.1101/2022.01.19.476913

**Authors:** Ida Calvi, Françoise Schwager, Monica Gotta

**Affiliations:** Department of Cell Physiology and Metabolism, Faculty of Medicine, University of Geneva, 1211 Geneva 4, Switzerland

## Abstract

Cell polarity relies on the asymmetric distribution of the conserved PAR proteins, which is regulated by phosphorylation/dephosphorylation reactions. While the kinases involved have been well studied, the role of phosphatases remains poorly understood. In *C. elegans* zygotes, phosphorylation of the posterior PAR-2 protein by the atypical protein kinase PKC-3 inhibits PAR-2 cortical localization. Polarity establishment depends on loading of PAR-2 at the posterior cortex. We show that the PP1 phosphatases GSP-1 and GSP-2 are required for polarity establishment in embryos. We find that co-depletion of GSP-1 and GSP-2 abrogates the cortical localization of PAR-2 and that GSP-1 and GSP-2 interact with PAR-2 via a PP1 docking motif in PAR-2. Mutating this motif *in vivo,* to prevent binding of PAR-2 to PP1, abolishes cortical localization of PAR-2, while optimizing this motif extends PAR-2 cortical localization. Our data suggest a model in which GSP-1/-2 counteract PKC-3 phosphorylation of PAR-2 allowing its cortical localization at the posterior and polarization of the one-cell embryo.

**SUMMARY:** Calvi et al. identify PP1 protein phosphatases as regulators of cell polarity in *C. elegans* embryos. Their results show that two redundant phosphatases, GSP-1 and GSP-2, interact with the polarity protein PAR-2 and control its localization and polarity establishment.

## INTRODUCTION

Cell polarity is a fundamental property of cells required for many aspects of cell and animal biology. In migrating cells, for example, a front-rear polarity regulates migration in response to chemokines or antigens (Llense & Etienne-Manneville, 2015); in stem cells, cell polarity is a pre-requisite for asymmetric cell division (Santoro et al., 2016); in epithelial cells, the apical-basal polarity axis is required to establish the barrier function of the epithelium (Riga et al., 2020; Rodriguez-Boulan & Macara, 2014;

Roignot et al., 2013).

In many different cells polarity is regulated by the conserved Partitioning defective (PAR) proteins, which have been identified in *C. elegans* (Goldstein & Macara, 2007; Rose & Gönczy, 2014).

The *C. elegans* zygote is a powerful model system for investigating the mechanism of cell polarity establishment and maintenance. The one-cell *C. elegans* embryo is polarized along the anterior-posterior (A-P) axis, with the anterior PAR proteins (the PDZ proteins PAR-3 and PAR-6, the atypical protein kinase C, PKC-3, and the small GTPase CDC-42, aPARs) enriched at the cortex in the anterior half of the embryo and the posterior PAR proteins (the ring finger protein PAR-2, the kinase PAR-1, the Lethal Giant Larvae orthologue, LGL-1, and the CDC-42 GAP CHIN-1, pPARs) enriched at the posterior cortex (reviewed in (Goehring, 2014; Lang & Munro, 2017)). This polarization results in a first asymmetric cell division, giving origin to two cells, AB and P1, with different size and fate. The zygote polarizes in two distinct phases, establishment and maintenance (Cuenca et al., 2003). Just after fertilization, PAR-3, PAR-6 and PKC-3 are uniformly distributed at the cortex, whereas PAR-1 and PAR-2 are in the cytoplasm (Fig. 1 A). Protein kinase C (PKC-3) phosphorylates PAR-2 and PAR-1 inhibiting their cortical localization and polarity establishment (Folkmann & Seydoux, 2019; Hao et al., 2006; Motegi et al., 2011). Polarity establishment relies on two redundant pathways, both dependent on the centrosomes. Shortly after the fertilization, a gradient of the mitotic kinase Aurora A (AIR-1) from the centrosomes of the paternal pronucleus triggers a cortical flow away from the newly defined posterior pole (Kapoor & Kotak, 2019; Klinkert et al., 2019; Reich et al., 2019; Zhao et al., 2019). This initiates the segregation of the aPARs to the anterior side of the embryo and liberates the posterior pole, allowing the localization of PAR-2 (and PAR-1) (Munro et al., 2004). A second, redundant pathway, relies on centrosomes-emanating microtubules, which promote loading of PAR-2 in the posterior (Fig. 1 A, (Motegi et al., 2011)). PAR-2 recruits PAR-1, thereby excluding the anterior PARs via PAR-1 dependent phosphorylation of PAR-3 (Motegi et al., 2011). Once polarity is established, a mutual antagonism between the anterior and posterior PAR proteins ensures polarity maintenance (reviewed in (Gubieda et al., 2020; Rose & Gönczy, 2014)).

**Figure 1:**
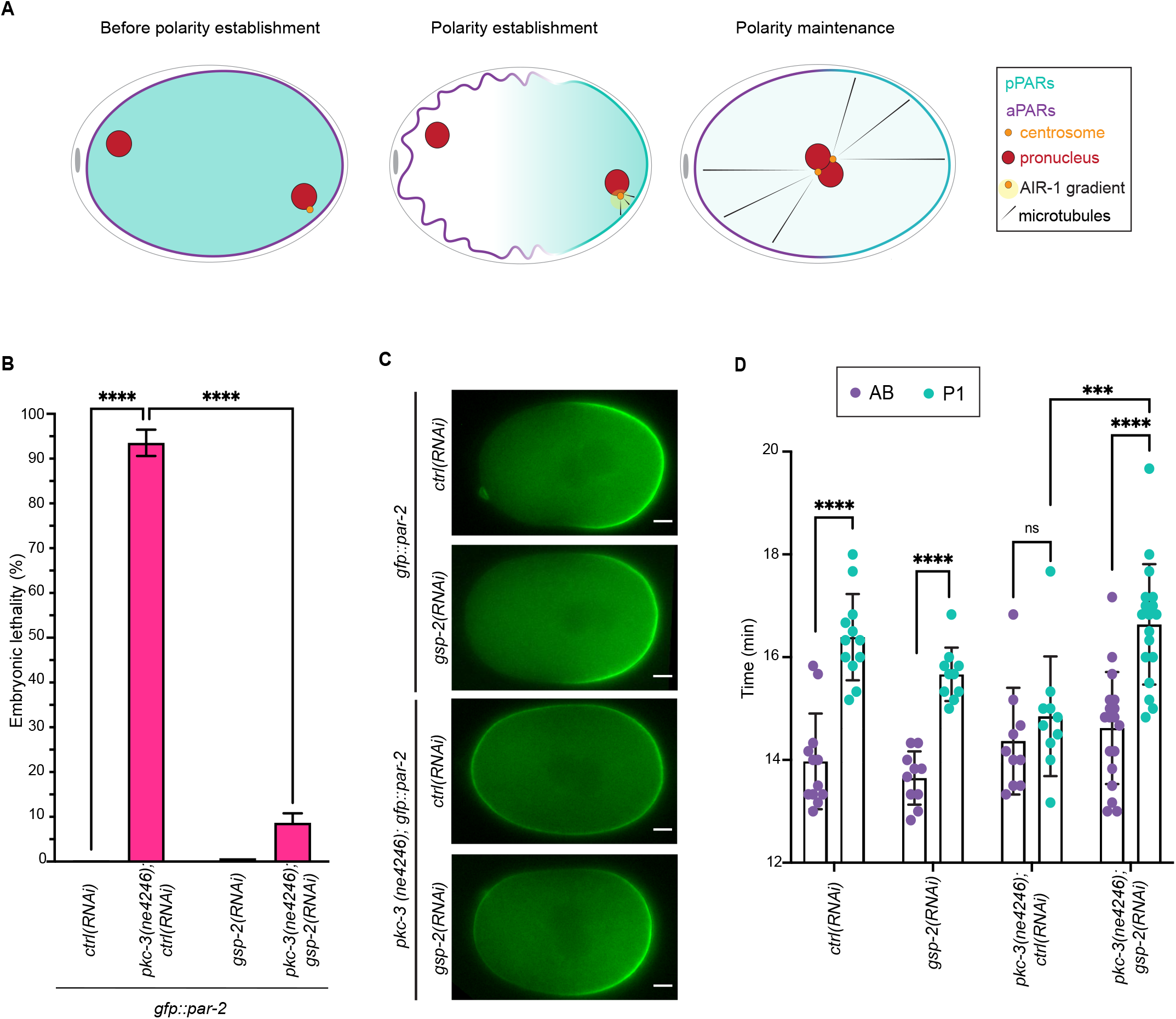
GSP-2 depletion suppresses *pkc-3(ne4246)* lethality and polarity defects. **(A)** Representation of the main phases that lead to zygote polarization. Left: early embryo before polarity establishment with anterior PARs uniformly at the cortex and posterior PARs in the cytoplasm. Middle: AIR-1 and microtubules dependent symmetry breaking. Right: polarity maintenance with anterior PARs at the anterior cortex and posterior PARs at the posterior cortex. **(B)** Embryonic lethality of *gfp::par-2* and *pkc-3(ne4246); gfp::par-2* embryos after *ctrl(RNAi)* and *gsp-2(RNAi).* The reported values correspond to the percentage of un-hatched embryos over the total progeny (larvae and un-hatched embryo). *gfp::par-*2; *ctrl(RNAi)*, n=3132, *gfp::par-2; gsp-2(RNAi),* n=2405, *pkc-3(ne4246); gfp::par-2; ctrl(RNAi),* n=1085 and *pkc-3(ne4246); gfp::par-2; gsp-2(RNAi*), n=1141. N=4. Mean is shown and error bars indicate SEM. The P-values were determined using 2way ANOVA “Tukey’s multiple comparisons test”. **(C)** Representative midsection frames of time-lapse imaging of embryos at the polarity maintenance stage: *gfp::par-*2; *ctrl(RNAi)*, n=11, *gfp::par-2; gsp-2(RNAi),* n=13, *pkc-3(ne4246); gfp::par-2; ctrl(RNAi),* n=15 and *pkc-3(ne4246); gfp::par-2; gsp-2(RNAi*), n=18, N=4. **(D)** Quantification of the cell cycle time: *ctrl(RNAi)*, n=12, *gsp-2(RNAi)*, n=10, *pkc-3(ne4246), ctrl(RNAi),* n=10, *pkc-3(ne4246); gsp-2(RNAi*), n=18. N=3. Mean is shown and error bars indicate SD. The P-values were determined using unpaired Student’s t-test between AB and P1 for each condition and between P1 *pkc-3(ne4246); ctrl(RNAi)* and P1 *pkc-3(ne4246); gsp-2(RNAi)*. ns p > 0.05, ***p<0.001, ****p < 0.0001. For all figures, the scale bars are 5 µm, anterior is to the left and posterior to the right.

PKC-3 phosphorylates PAR-2 and inhibits its membrane association (Hao et al., 2006). Despite the fact that centrosomal microtubules can protect PAR-2 from PKC-3 phosphorylation and therefore promote PAR-2 membrane association, mutations of the PAR-2 microtubule binding sites delay but do not abolish posterior cortical loading of PAR-2 in presence of normal cortical flows (Motegi et al., 2011). On the contrary, when the PKC-3 phosphorylation sites in PAR-2 are mutated to mimic phosphorylation, PAR-2 localization at the posterior cortex is abrogated, resulting in a defect in polarity establishment (Hao et al., 2006). This suggests that PAR-2 phosphorylation by PKC-3 must be relieved to ensure PAR-2 posterior cortical localization and hence proper polarity establishment.

Here we show that the PP1 phosphatase GSP-2 plays an important role in polarity establishment. GSP-2 depletion suppresses the lethality (as previously shown (Fievet et al., 2013)) and the polarity defects of a temperature sensitive *pkc-3* mutant, suggesting that GSP-2 antagonizes PKC-3 function. GSP-2 depletion also results in defects in PAR-2 cortical localization that are exacerbated by the depletion of the redundant PP1 phosphatase GSP-1. Consistent with a role of GSP-2 and GSP-1 in PAR-2 dephosphorylation, PAR-2 contains a PP1 binding motif, which we show to be required for the interaction with GSP-1 and GSP-2 in two-hybrid assays. Mutations of this site known to abolish PP1 binding (Meiselbach et al., 2006) abrogate PAR-2 cortical localization and polarity establishment *in vivo* (this study). On the opposite, mutations that optimize the PP1 binding motif, extend PAR-2 cortical localization (Meiselbach et al., 2006).

Our work identifies the PP1 phosphatases GSP-2 and GSP-1 as critical regulators of PAR-2 cortical localization and polarity establishment in the *C. elegans* embryo.

## RESULTS

### Depletion of GSP-2 suppresses the lethality and polarity phenotypes of a temperature sensitive *pkc-3* mutant

GSP-2 has been identified as a suppressor of the embryonic lethality caused by a temperature sensitive mutant allele of *pkc-3* (*pkc-3(ne4246)* (Fievet et al., 2013)). PKC-3 phosphorylates PAR-2 and inhibits its localization at the anterior cortex (Hao et al., 2006; Motegi et al., 2011). In the *pkc-3(ne4246)* strain, at restrictive temperature, PAR-2 occupies the entire cortex (Rodriguez et al., 2017) resulting in polarity defects and embryonic lethality. We therefore asked whether depletion of GSP-2 was able to rescue the aberrant localization of PAR-2 observed in the *pkc-3(ne4246)* embryos. Embryonic lethality of *the pkc-3(ne4246); gfp::par-2*; *ctrl(RNAi)* strain at 24°C (see Material and Methods) was about 93.5% (Fig. 1 B). Consistent with the high embryonic lethality, embryos from *pkc-3(ne4246); gfp::par-2; ctrl(RNAi)* mutant strain had impaired A-P polarity, with PAR-2 not being restricted anymore to the posterior, but distributed all around the cortex in one-cell stage embryos and partitioned symmetrically into the daughter AB and P1 cells (Fig. 1 C and Video 1). Depletion of GSP-2 in *gfp::par-2* worms did not result in embryonic lethality, and PAR-2 was localized at the posterior cortex (Figs. 1 B, C). Depletion of GSP-2 in the *pkc-3(ne4246); gfp::par-2* mutant rescued the embryonic lethality (Fig. 1 B). We found that PAR-2 localization was restored at the posterior cortex in one-cell stage embryos and was restricted to the P1 blastomere in two-cells stage embryos (Fig. 1 C and Video 1). Therefore, depletion of GSP-2 in the *pkc-3(ne4246)*; *gfp::par-2* rescued the embryonic lethality and PAR-2 localization defects of early embryos.

Polarity controls the posterior positioning of the mitotic spindle leading to a more posterior cleavage and to two-cell embryos with a bigger anterior cell (AB) and a smaller posterior cell (P1). Depletion of GSP-2 in the *gfp::par-2* control strain resulted in a higher AB/P1 ratio compared to *ctrl(RNAi)* (Fig. S1 A). In the *pkc-3(ne4246); gfp::par-2; ctrl(RNAi)* the cleavage furrow was more shifted towards the anterior resulting in a reduced size asymmetry of the daughter cells. This phenotype was also rescued by depletion of GSP-2 (Fig. S1 A).

*C. elegans* two-cell embryos divide asynchronously, with the anterior AB cell dividing about 2 minutes before P1. This asynchrony is lost in the *pkc-3* mutant. We therefore tested whether GSP-2 depletion was able to rescue this phenotype. AB divided roughly 2 minutes before P1 both in *ctrl(RNAi)* and in *gsp-2(RNAi)* embryos, whereas in the *pkc-3(ne4246)* mutant allele, the division time of AB and P1 was not significantly different (Fig. 1 D and Video 2). Depletion of GSP-2 in the *pkc-3(ne4246)* strain was able to restore the asynchrony between AB and P1 (Fig. 1 D and Video 2).

One mechanism behind the regulation of asynchrony is the polarity dependent localization of the mitotic kinase Polo-Like Kinase 1 (PLK-1). In the one cell embryo, PLK-1 becomes enriched in the anterior cytoplasm and, at division, it is preferentially segregated in AB. This enrichment in AB triggers the earlier division of this blastomere (Budirahardja & Gönczy, 2008; Nishi et al., 2008; Rivers et al., 2008). We therefore investigated the localization of PLK-1. In *gsp-2(RNAi)* embryos, PLK-1 was enriched in the anterior AB cell as in *ctrl(RNAi)* embryos (Fig. S1 B). In the *pkc-3(ne4246)* embryos, the AB/P1 ratio of PLK-1 levels was close to 1, consistent with what has been previously reported in absence of the aPARs (Budirahardja & Gönczy, 2008; Nishi et al., 2008; Rivers et al., 2008). In *pkc-3(ne4246); gsp-2(RNAi)* embryos PLK-1 anterior enrichment was restored (Fig. S1 B).

Taken together, these results show that depletion of GSP-2 suppresses the embryonic lethality and the polarity defects of the *pkc-3(ne4246)* mutant allele.

### GSP-2 antagonizes PKC-3 in the regulation of PAR-2 localization at the posterior cortex

The results that GSP-2 depletion rescues the PAR-2 localization defects of *pkc-3(ne4246)* mutant embryos suggest that GSP-2 and PKC-3 antagonize each other in the regulation of PAR-2 cortical localization. PKC-3 phosphorylates PAR-2 thereby inhibiting its membrane localization (Hao et al., 2006). Since GSP-2 is a phosphatase, one possibility is that GSP-2 dephosphorylates PAR-2 allowing its posterior cortical localization. If this was the case, depletion of GSP-2 in control embryos should result in a reduction of cortical PAR-2, as PAR-2 would not be dephosphorylated. When GSP-2 was depleted in *gfp::par-2* embryos, PAR-2 was localized at the posterior cortex (Fig. 1 C and 2 A). However, the size of the PAR-2 domain was smaller (Fig. 2 A), indicating that GSP-2 contributes to the formation of a PAR-2 domain of the correct size.

**Figure 2:**
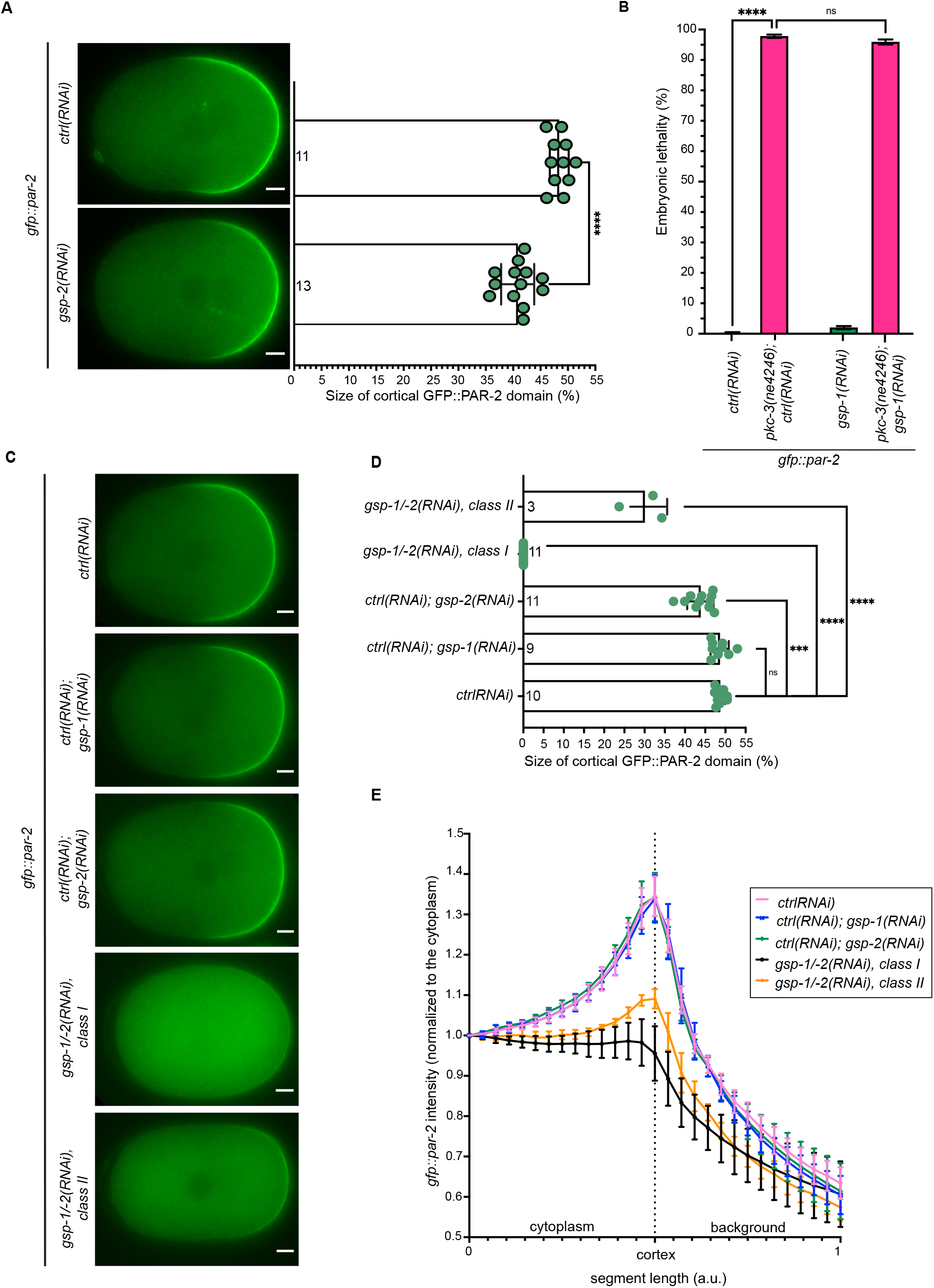
Co-depletion of GSP-1 and GSP-2 interferes with PAR-2 cortical posterior localization. **(A)** Left: representative midsection frames of time-lapse imaging of *gfp::par-2* zygotes at pronuclear meeting, comparing *ctrl(RNAi)* and *gsp-2(RNAi)*. N=4. Right: quantification of the *gfp::par-2* size domain in live zygotes at pronuclear meeting. Numbers inside the bars indicate sample size. N=4. Mean is shown and error bars indicate SD. The P-values were determined using unpaired Student’s t-test. **(B)** Embryonic lethality of *gfp::par-2* and *pkc-3(ne4246); gfp::par-2* embryos after *ctrl(RNAi)* and *gsp-1(RNAi).* The reported values correspond to the percentage of un-hatched embryos over the total progeny (larvae and un-hatched embryo). *gfp::par-*2; *ctrl(RNAi)*, n=2040, *gfp::par-2; gsp-1(RNAi),* n=1586, *pkc-3(ne4246); gfp::par-2; ctrl(RNAi),* n=751 and *pkc-3(ne4246); gfp::par-2; gsp-1(RNAi*), n=382. N=3. Mean is shown and error bars indicate SEM. The P-values were determined using 2way ANOVA “Tukey’s multiple comparisons test”. **(C)** Representative midsection frames of time-lapse imaging of *gfp::par-2* zygotes at pronuclear meeting, comparing *ctrl(RNAi)*, n= 10, *ctrl(RNAi); gsp-1(RNAi)*, n= 9, *ctrl(RNAi); gsp-2(RNAi),* n= 11 and *gsp-1/- 2(RNAi) class I,* n= 11 and *class I,I* n=3. N=3. Zygotes co-depleted of both GSP-1 and GSP-2 show PAR-2 uniformly in the cytoplasm in most of the embryos (class I, 78.6%); class II embryos (21.4%) show a weak PAR-2 localization at the cortex compared to *ctrl(RNAi)*. **(D)** Quantification of the *gfp::par-2* size domain in live zygotes at pronuclear meeting. Numbers inside the bars indicate sample size. N=3. Mean is shown and error bars indicate SD. The P-values were determined using unpaired Student’s t-test. **(E)** PAR-2 line profile of live zygotes at pronuclear meeting comparing *ctrl(RNAi)*, n=10, *ctrl(RNAi); gsp-1 (RNAi)*, n=9 *ctrl(RNAi); gsp-2(RNAi)* n=11 and *gsp-1/-2(RNAi) class I,* n=11 and *class II,* n=3. Mean is shown and error bars indicate SD. N=3. ns p > 0.05, ***p<0.001, ****p < 0.0001.

*C. elegans* embryos express a second PP1 catalytic subunit, GSP-1, which is 85% identical to GSP-2 in the amino acid sequence (Sassa et al., 2003). GSP-1 and GSP-2 have the same localization in embryos, and they are functionally redundant (Mangal et al., 2018; Peel et al., 2017). We therefore investigated whether GSP-1 and GSP-2 are redundant in promoting PAR-2 cortical localization in one-cell embryos. We first asked whether GSP-1 depletion rescued the embryonic lethality of *pkc-3(ne4246); gfp::par-2* worms. Depletion of GSP-1 in *gfp::par-2* worms did not result in embryonic lethality, and its depletion in the *pkc-3(ne4246)*; *gfp::par-2* mutant did not rescue the embryonic lethality of this strain (Fig. 2 B). Consistent with this, PAR-2 was detected uniformly at the cortex in both *pkc-3(ne4246)*; *gfp::par-2*; *ctrl(RNAi)* and *pkc-3(ne4246)*; *gfp::par-2, gsp-1(RNAi)* (Fig. S2 A and Video 3). In addition, embryos depleted of GSP-1 did not show a smaller PAR-2 domain compared to *ctrl(RNAi)* embryos (Fig. S2 B). Therefore, in contrast to GSP-2 depletion, GSP-1 depletion did not suppress the embryonic lethality and polarity defects observed in the *pkc-3(ne4246*) mutant allele and did not impair the size of the PAR-2 domain.

We then asked if co-depletion of GSP-1 and GSP-2 in the *gfp::par-2* embryos impaired PAR-2 localization. Consistent with the previous results, we could observe a small but significant reduction of the size of the PAR-2 domain in the *ctrl(RNAi); gsp-2(RNAi)* embryos, while the PAR-2 domain was not reduced in the *ctrl(RNAi); gsp-1(RNAi)* embryos (Fig. 2 C, D). The intensity of the PAR-2 domain did not change in the single depletion (Fig. 2 E). In the GSP-1/-2 depleted embryos we observed two phenotypes: in the majority of the embryos PAR-2 was mostly cytoplasmic (*class I*, 78.6%), as shown by an almost flat intensity profile, whereas a smaller percentage of embryos (*class II*, 21.4%) showed a weak localization of PAR-2 at the posterior cortex (Figs. 2 C, D, E and Video 4).

To conclude, co-depletion of GSP-1 and GSP-2 results in a defect in PAR-2 posterior cortical localization. In addition, our data suggest that GSP-2 has a leading role in the regulation of PAR-2 localization but GSP-1 can compensate in the absence of GSP-2.

### GSP-1 and GSP-2 interact with PAR-2

GSP-2 and a PAR-2 N-terminal fragment (Fig. 3 A) were identified as interactors in large scale two-hybrid screen (Koorman et al., 2016). The genetic and physical interactions suggest that GSP-2 (redundantly with GSP-1) is the phosphatase that antagonizes PKC-3 phosphorylation of PAR-2. We therefore tested whether GSP-2 and GSP-1 interacted with PAR-2. As previously shown (Koorman et al., 2016), we confirmed that GSP-2 interacted with PAR-2 (Fig. 3 B). Consistent with the redundancy in regulating PAR-2 localization, GSP-1 also interacted with PAR-2 (Fig. 3 B).

**Figure 3:**
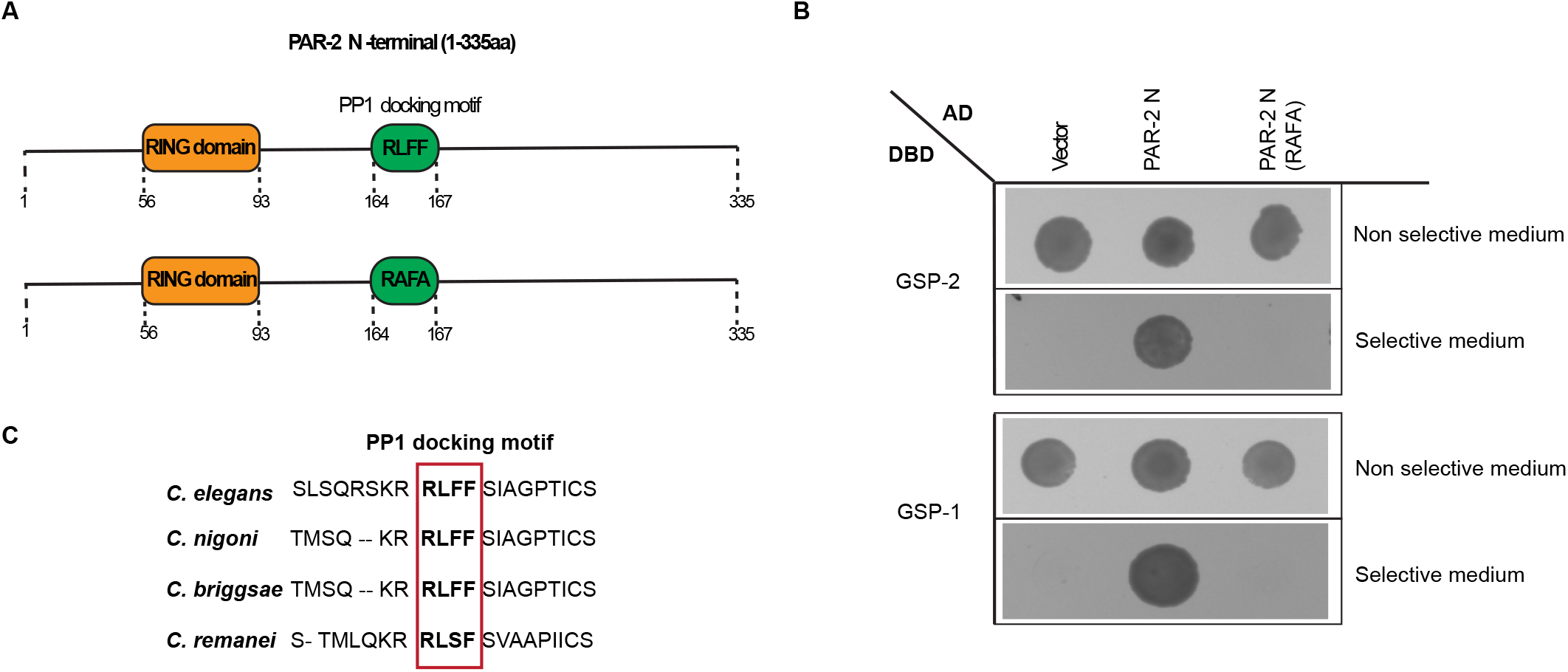
PAR-2 interacts with both GSP-1 and GSP-2. **(A)** Schematic representation of the N-terminal (1-335aa) of both *wild-type* PAR-2 (upper panel) and PP1 mutant (RAFA, lower panel). **(B)** Yeast two-hybrid assay showing the interaction between PAR-2 N (wild type and the RAFA mutant) and GSP-2 and GSP-1. Yeasts were transformed with the indicated plasmids and were grown on selective and non-selective medium. Interaction between bait and prey results in growth on selective medium (50mM 3AT). **(C)** Alignment of the PP1 motif in PAR-2 for *C. elegans*, *C. nigoni*, *C. briggsae* and *C. remanei*. Figure re-elaborated from alignment obtained with Blast (https://blast.ncbi.nlm.nih.gov ).

Analysis of the PAR-2 amino acid sequence revealed the presence of a degenerate PP1 docking motif (164 RLFF 167) in the N-terminus of PAR-2, conserved in closely related nematode species (Fig. 3 A, upper panel, and Fig. 3 C). This suggests that PAR-2 may physically interact with PP1 through this motif (Egloff et al., 1997; Hendrickx et al., 2009; Wakula et al., 2003; Zhao & Lee, 1997). To assess if the PP1 docking motif present in PAR-2 is required for the interaction with GSP-1 and GSP-2, we mutated two critical amino acid residues into Alanine (referred as PAR-2 (RAFA), Fig. 3 A, lower panel). These substitutions have previously been shown to interfere with the binding between PP1 phosphatases and their substrates (Meiselbach et al., 2006; Moreira et al., 2019). Interestingly, neither GSP-1 nor GSP-2 could interact with PAR-2 in the Yeast Two Hybrid system when the PP1 docking motif in PAR-2 was mutated (Fig. 3 B).

These data show that PAR-2 interacts with both GSP-2 and GSP-1 in the two-hybrid assay, and this interaction depends on a PP1 docking motif in PAR-2.

### Mutations in the PP1 docking motif of PAR-2 result in polarity defects

We set out to assess whether cell polarity was impaired if the PP1 docking motif in PAR-2 was mutated. We generated a strain in which the PP1 RLFF motif was mutated to RAFA in the endogenous *gfp::par-2* (referred as *gfp::par-2(RAFA)*). The homozygote mutant strain exhibited high embryonic lethality (95.4% ± 4.5 SEM) and the surviving progeny was sterile (Fig. S3 A). Our control, referred to as *gfp::par-2°*, was a mixture of homozygous worms expressing wild-type *gfp::par-2* from both alleles and heterozygous worms expressing wild-type *gfp::par-2* from one allele and mutant *gfp::par-2(RAFA)* from the other allele. We could not detect any difference in cortical PAR-2 domain size and intensity in the *gfp::par-2°* population compared to the *gfp::par-2* (Figs. S3 B, C).

The *gfp::par-2(RAFA)* mutant displayed phenotypes consistent with the impairment of polarity establishment. PAR-2 remained mostly cytoplasmic (Figs. 4 A, B and Video 5), consistent with the phenotype of embryos co-depleted of both PP1 catalytic subunits and similar to what has been shown with the phosphomimetic mutant of PAR-2, in which 7 PKC-3 phosphorylation sites have been mutated to glutamic acid (Hao et al., 2006). The position of the cleavage furrow during the first cell division was variable and shifted towards the anterior, resulting in a more symmetric first cell division with AB and P1 of equal size (Fig. S3 D).

**Figure 4:**
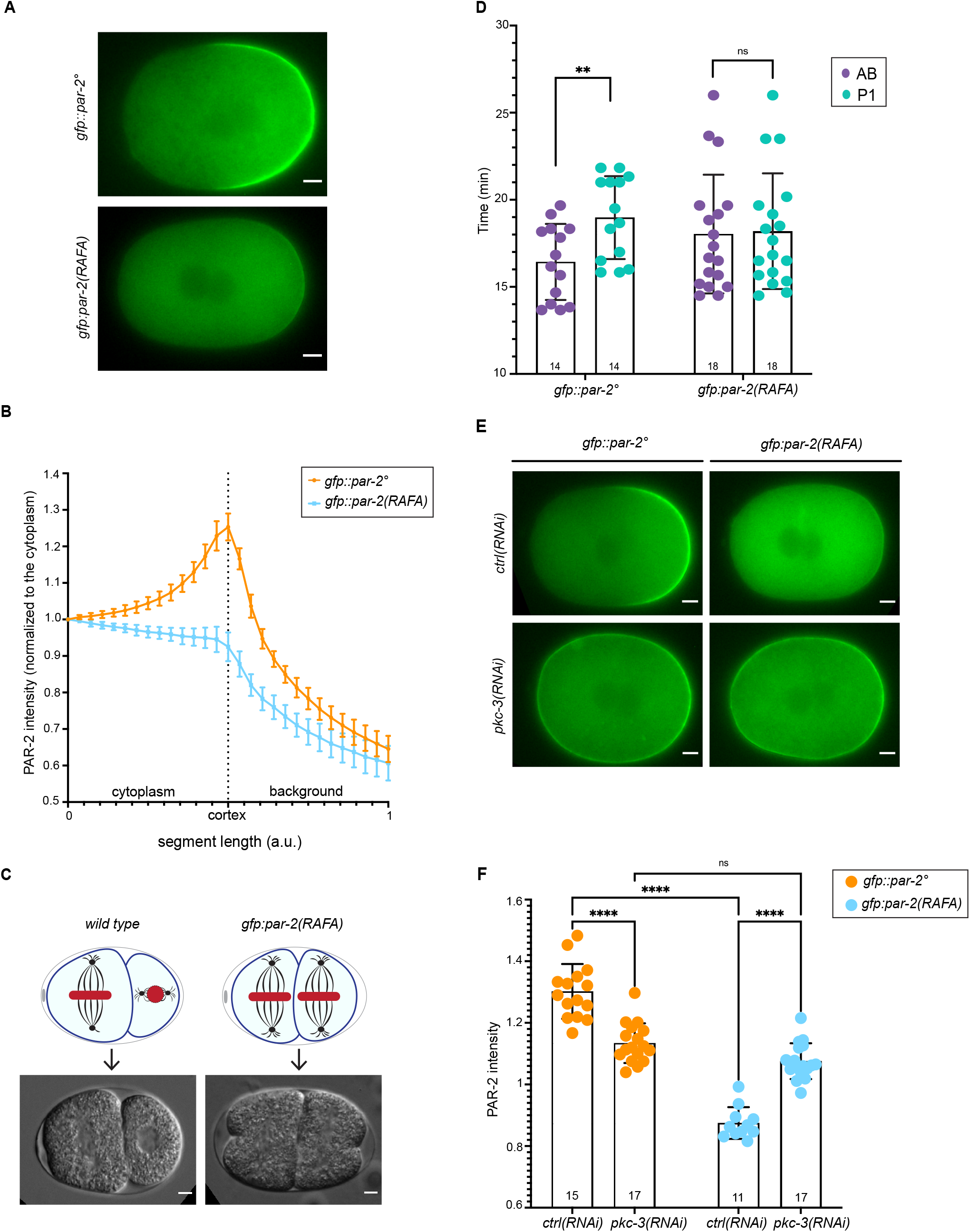
The PP1 binding site in PAR-2 is required *in vivo* for PAR-2 cortical posterior localization. **(A)** Representative midsection frames of time-lapse imaging of *gfp::par-2°* (the symbol ° indicates a mixture between wild-type *gfp::par-2* homozygous worms expressing wild-type *gfp::par-2* and heterozygous worms expressing wild type *gfp::par-2* from one allele and mutant *gfp::par-2(RAFA)* from the other allele) and *gfp::par-2(RAFA)* embryos at pronuclear meeting (n=16 and n=17 respectively) N=6. **(B)** PAR-2 line profile of live zygotes at pronuclear meeting: *gfp::par-2°*, n= 16, and *gfp::par-2(RAFA),* n=17, N=6. Mean is shown and error bars indicate SD. **(C)** *wild-type* and *gfp::par-2(RAFA)* dividing two-cell stage embryos. The upper panel show a schematic representation, whereas in the lower panel images from DIC time-lapse movies are shown. In the schematics, the nucleus and the metaphase plate are in red, black lines are microtubules, black circles are centrosomes. **(D)** Quantification of the cell cycle time in *gfp::par-2°* and and *gfp::par-2(RAFA)* embryos. Numbers inside the bars indicate sample size. N=6. Mean is shown and error bars indicate SD. The P-values were determined using Student’s t-test. **(E)** Representative midsection frames of time-lapse imaging of *gfp::par-2°* and *gfp::par-2(RAFA)* one-cell embryos at pronuclear meeting. *gfp::par-2°*; *ctrl(RNAi)* n= 15, *pkc-3(RNAi); gfp::par-2°* n= 17, *gfp::par-2(RAFA); ctrl(RNAi)* n= 11 and *pkc-3(RNAi); gfp::par-2(RAFA)* n=17, N=7. **(F)** Quantification of PAR-2 intensity obtained as a ratio of PAR-2 posterior cortex intensity over the posterior cytoplasm. Numbers inside the bars indicate sample size. N=7. Mean is shown and error bars indicate SD. The P-values were determined using 2way ANOVA “Tukey’s multiple comparisons test”. ns p > 0.05, **p < 0.01, ****p < 0.0001.

In control embryos, the P1 spindle rotates in order to be oriented along the anterior-posterior (A-P) axis, whereas the AB spindle is orthogonal to the A-P axis (Fig. 4 C). Mutations that interfere with the establishment of polarity can alter spindle orientation at the second division (Cheng et al., 1995; Kemphues et al., 1988). We found that in the *gfp::par-2(RAFA)* mutant embryos, the P1 spindle failed to rotate, resulting in an irregular arrangement of cells at the four-cell stage (Fig. 4 C). Furthermore, the AB and P1 blastomeres divided synchronously (Fig. 4 D and Video 6). Therefore, mutation of the PP1 docking motif in the endogenous PAR-2 strongly reduced PAR-2 cortical localization and led to polarity defects similar to the ones observed in *par-2* loss of function embryos (Cheng et al., 1995; Kemphues et al., 1988).

If the phosphatase PP1 antagonizes the kinase activity of PKC-3 on PAR-2, depletion of PKC-3 in the *gfp::par-2(RAFA)* mutant strain should result in PAR-2 localizing around the entire cortex. *pkc-3(RNAi); gfp::par-2(RAFA)* embryos displayed uniform PAR-2 cortical localization in all the embryos analyzed (Figs. 4 E, F).

These results suggest that the interaction between PAR-2 and GSP-1/-2 has a crucial role in the regulation of PAR-2 cortical localization and establishment of polarity.

### Optimizing the PP1 binding motif of PAR-2 results in PAR-2 aberrant cortical localization

One of the hallmarks of the PP1 docking motif is the high degeneracy at key positions of the motif (Davey et al., 2015). The optimal PP1 docking motif is the RVxF sequence (Wakula et al., 2003) and the Valine was shown *in vitro* to contribute to a stronger interaction between PP1 and its substrates (Meiselbach et al., 2006). The PP1 docking motif in PAR-2 presents a Leucine in position 165 instead of the optimal Valine (Fig. 3 A). We therefore asked if optimizing the PP1 docking motif by mutating the Leucine to Valine *in vivo* would alter the cortical localization of PAR-2. We hypothesized that if this mutation improves the binding of PAR-2 to PP1 phosphatases, PAR-2 cortical domain may be extended.

We generated a strain where the Leucine 165 in the PP1 docking motif was mutated to Valine in the endogenous *gfp::par-2* (referred as *gfp::par-2(L165V)*). The mutant strain did not show embryonic lethality compared to *gfp::par-2* worms (0.31% ± 0.25 SEM vs 0.17% ± 0.06 SEM respectively) and the progeny was fertile. All *gfp::par-2(L165V)* zygotes, analyzed before symmetry breaking, displayed PAR-2 localization uniformly around the entire cortex, a phenotype that was not observed in control embryos (Fig. 5 A). During pronuclei migration we detected both an anterior and a posterior PAR-2 domain in all the zygotes analyzed, indicating that in this strain restriction of PAR-2 to the posterior cortex is impaired. The anterior PAR-2 domain remained in 32% of embryos in mitosis and 16% of two-cell embryos (Fig. 5 A and Video 7).

**Figure 5:**
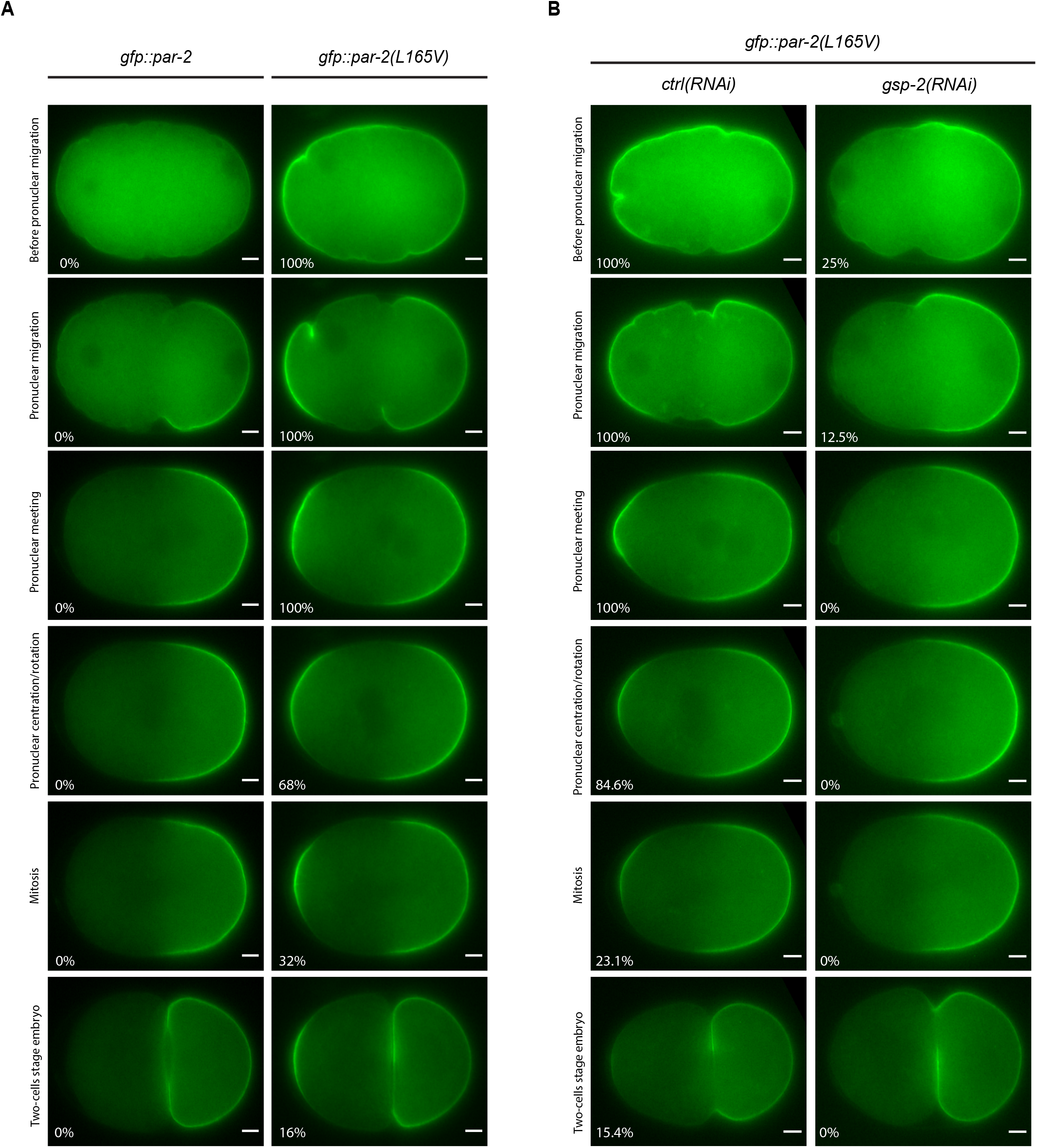
The L165V mutation in PAR-2 causes atypical PAR-2 cortical localization. **(A)** Representative midsection frames of time-lapse videos of *gfp::par-*2 and *gfp::par-2(L165V)* embryos at different cell division stages (for embryos before pronuclear migration: n= 9 and n=14 respectively, for the other embryos stage showed n=14 and n=25 respectively). Percentage on the bottom left of each image indicates the percentage of embryos with an anterior PAR-2 domain, except for the top panel (before pronuclear migration) in which indicates the percentage of embryos with PAR-2 around the entire cortex. N=4. **(B)** Representative midsection frames of time-lapse imaging of *gfp::par-*2 and *gfp::par-2(L165V)* one-cell embryos at different cell division stages, *ctrl(RNAi),* n=13 and *gsp-2(RNAi)*, n=16. For the top panel (before pronuclear migration) n= 6 and n=8 for *ctrl(RNAi)* and *gsp-2(RNAi)* respectively. N=3. As in panel A, percentage on the bottom left of each embryo indicates the percentage of embryos with an anterior PAR-2 domain, with the exception of the top panel.

If the rate of PP1 dependent dephosphorylation was increased in the *gfp::par-2(L165V)* mutant strain, depletion of GSP-2 should rescue the aberrant localization of PAR-2 in this mutant. PAR-2 showed normal localization at the posterior pole in 75% of the *gfp::par-2(L165V); gsp-2(RNAi)* early embryos and in 87.5% of the embryos analyzed at the onset of pronuclear migration; from pronuclear migration to later stage all the embryos analyzed showed only one PAR-2 domain at the posterior cortex (Fig. 5 B).

Collectively these results show that optimizing *in vivo* the PP1 binding motif of PAR-2 results in aberrant cortical localization of PAR-2 before and during polarity establishment. This localization is corrected in most but not all embryos at later stages, suggesting that other yet unknown mechanisms are involved in the restriction of PAR-2 at the posterior cortex.

### Depletion of GSP-2 in a temperature sensitive *plk-1* mutant allele rescues aberrant PAR-2 localization

An anterior PAR-2 domain similar to the one observed in the *gfp::par-2(L165V)* mutant has been reported in embryos in meiosis (Wallenfang & Seydoux, 2000) and in embryos where the activity of the mitotic kinases PLK-1 and Aurora A (AIR-1) have been reduced (Kapoor & Kotak, 2019; Klinkert et al., 2019; Noatynska et al., 2010; Reich et al., 2019; Schumacher et al., 1998). We wondered whether PLK-1, which is enriched in the anterior cytoplasm of the embryo, might contribute to restrict the activity of the phosphatases at the posterior. To test this, we used the temperature sensitive mutant allele *plk-1(or683)*. If the anterior PAR-2 domain observed in the *plk-1(or683)* mutant allele is due to the increased PP1 activity, the depletion of GSP-2 in the *plk-1(or683)* mutant allele should reduce the number of embryos showing this phenotype.

We stained *plk-1(or683); ctrl(RNAi)* embryos with PAR-2 antibodies and found that PAR-2 localized at both the anterior and posterior domains, whereas in the *wild type; ctrl(RNAi)* embryos only one PAR-2 domain was detected (Fig. 6). Depletion of GSP-2 in the *plk-1(or683)* embryos resulted in a significant reduction of embryos with the anterior PAR-2 domain (Fig. 6).

**Figure 6:**
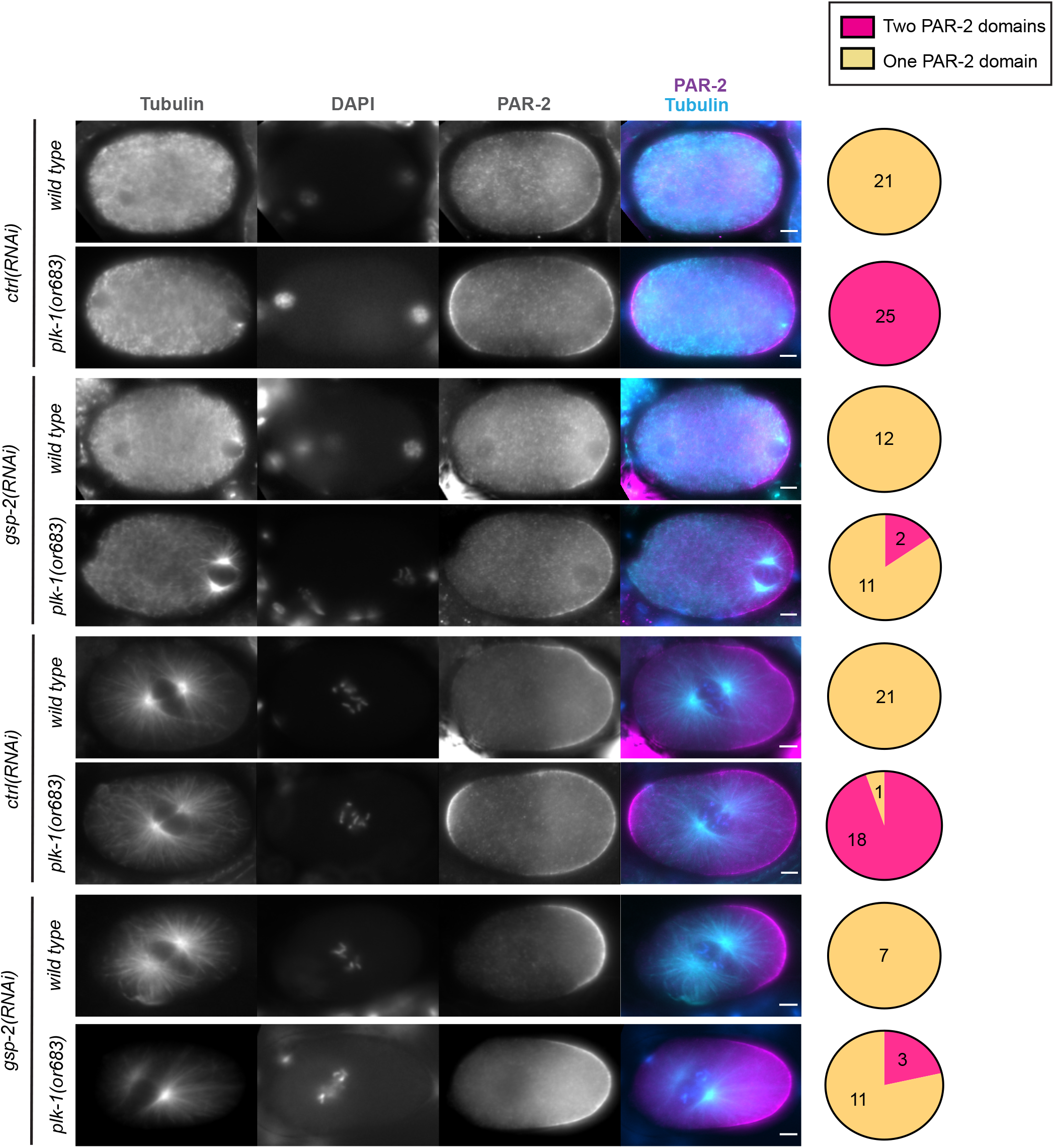
PLK-1 limits the activity of GSP-2. Representative images of fixed embryos of the indicated genotypes during pronuclei migration and at pronuclei centration/rotation, stained with anti-tubulin and anti-PAR-2 antibodies. Tubulin is in cyan, PAR-2 in magenta. The number of embryos analyzed is shown on the right in the pie-chart graph. N=5.

These data suggest that the anterior PAR-2 domain in the *plk-1(or683)* mutant allele depends on the GSP-2 phosphatase, and that PLK-1 and GSP-2 antagonizes each other in the regulation of cortical PAR-2 at the anterior.

## Discussion

Establishment of the anterior-posterior axis is an essential process to ensure asymmetric cell division and proper development of the *C. elegans* embryo. Here we show that in the one-cell embryo this process is regulated by PP1 phosphatases. We find that the PP1 phosphatase GSP-2 and the redundant GSP-1 are required for the loading of PAR-2 at the cortex. Mutations in the PP1 binding motif of PAR-2, which abrogate the interaction with GSP-1 and GSP-2 in yeast two-hybrid assays, result, *in vivo*, in the inability of PAR-2 to properly localize at the cortex. Our data suggest a model in which GSP-2 and GSP-1 counterbalance the activity of PKC-3 in the early embryo, allowing PAR-2 posterior cortical localization.

These data also support previous findings in the field. Establishment of cell polarity in the one-cell *C. elegans* embryos relies on cortical flows that displace aPARs, including PKC-3, from the posterior allowing loading of PAR-2 and PAR-1 (Munro et al., 2004; Shelton et al., 1999). In addition to this pathway, binding of PAR-2 to astral microtubules protects PAR-2 from PKC-3 phosphorylation and promotes its posterior cortical localization, therefore contributing to polarity establishment (Motegi et al., 2011). However, abolishing binding of PAR-2 to microtubules does not abrogate polarity establishment in the presence of cortical flows while a PAR-2 mutant that mimics PKC-3 phosphorylation is unable to polarize the embryo (Hao et al., 2006; Motegi et al., 2011). This suggests that the major polarization pathway involving the flows needs the action of phosphatases to dephosphorylate PAR-2, as shown by our data.

The PP1 phosphatase GSP-2 was previously shown to be a suppressor of the embryonic lethality of the *pkc-3(ne4246)* mutant allele (Fievet et al., 2013). We find that depletion of GSP-2 in the *pkc-3(ne4246)* allele also rescues the polarity-related defects observed in the *pkc-3(ne4246)* mutant alone. Depletion of GSP-2 in otherwise control strains did not result in major defects in polarity establishment, as it would have been expected if GSP-2 was the only phosphatase targeting PAR-2. Co-depletion of the redundant GSP-1 does result in polarity establishment defects. However, depletion of GSP-1 alone was not able to rescue the lethality of *pkc-3(ne4246).* Although the polarity phenotype is enhanced in absence of both GSP-1 and GSP-2, only GSP-2 depletion can rescue *pkc-3(ne4246)* embryonic lethality and polarity defects and result in a smaller size of the PAR-2 domain in one-cell embryos, suggesting that GSP-2 is the phosphatase that has a more important role in polarity regulation. This is reminiscent of previous work showing that GSP-1 and GSP-2 have a redundant role in the regulation of centriole amplification. Interestingly, in this process, GSP-1 plays a more important role (Peel et al., 2017).

We find that PAR-2 contains a PP1 docking motif, a common motif used by PP1 phosphatases to physically interact with their substrates (Egloff et al., 1997; Hendrickx et al., 2009; Wakula et al., 2003; Zhao & Lee, 1997). This motif is also present in PAR-2 of closely related nematode species, suggesting that dephosphorylation of PAR-2 by PP1 is a conserved process in nematodes. In this context it is interesting to note that in *C. elegans* embryos the ortholog of the lethal giant larvae protein, LGL-1, can partially compensate for the loss of PAR-2 (Beatty et al., 2010; Hoege et al., 2010). In flies, Lgl is also an important regulator of polarity (Su et al., 2012). In epithelial cells, Lgl is removed from the cortex during mitosis in an aPKC and Aurora A phosphorylation dependent manner. Restoration of cortical localization of Lgl is essential to maintain cell polarity and tissue architecture and is regulated by a PP1 phosphatase (Moreira et al., 2019), similar to what we observe for PAR-2 in *C. elegans* zygotes.

Consistent with the role of the PP1 docking motif for the interaction with PAR-2 and the phenotype observed with the co-depletion of both GSP-1 and GSP-2, mutations of the two important amino acids in the PP1 docking motif (Leucine165 to Alanine and Phenilalanine167 to Alanine, referred to in the text as *gfp::par-2(RAFA)*) abrogate the interaction with PAR-2 in the yeast two-hybrid system; more importantly, *in vivo,* these mutations impair the localization of PAR-2 at the posterior cortex and embryo viability, resembling the *par-2* loss-of-function phenotype, consistent with the fact that dephosphorylation of PAR-2 is required for polarity establishment (Hao et al., 2006) (Figs. 4 and S3). The *gfp::par-2(RAFA)* mutant showed two interesting features. First, heterozygote embryos did not show any difference in PAR-2 cortical levels and PAR-2 domain size compared to homozygote wild type worms. We speculate that thanks to the property of PAR-2 to form oligomers (Arata et al., 2016), mutant PAR-2 can form oligomers with wild type PAR-2 and localize properly to the cortex. Second, we also noticed that the *gfp:par-2(RAFA)* mutant shows a weak PAR-2 localization at the posterior cortex (Video 5). One possibility is that in this mutant the microtubule redundant pathway protects some PAR-2 from phosphorylation by PKC-3, but this is not sufficient alone to ensure proper establishment of polarity.

Based on previous *in vitro* studies (Meiselbach et al., 2006) we have also optimized *in vivo* the PP1 binding motif. The Leucine165 to Valine mutant (referred to as *gfp::par-2(L165V)*) showed an aberrant PAR-2 localization around the entire cortex prior to polarity establishment. During polarity establishment and at later stages, some of the embryos displayed an anterior and a posterior PAR-2 domain. This phenotype is reminiscent of the phenotype observed in embryos where PLK-1 and AIR-1 have been depleted (Kapoor & Kotak, 2019; Klinkert et al., 2019; Noatynska et al., 2010; Reich et al., 2019; Schumacher et al., 1998). Aurora A can be an activator of Polo like kinase (Macůrek et al., 2008; Seki et al., 2008; Tavernier et al., 2015). PLK-1 in *C. elegans* embryos is enriched in the anterior cytoplasm (Budirahardja & Gönczy, 2008; Nishi et al., 2008; Rivers et al., 2008). One possibility is that Aurora A activated-PLK-1 contributes to keep GSP-2 activity low at the anterior, so that PAR-2 remains cytoplasmic (see Fig. 7). This is consistent with previous genetic data showing that PLK-1 depletion can rescue PAR-2 cortical localization in *par-2* temperature sensitive mutants (Noatynska et al., 2010) and our current data that GSP-2 depletion in *plk-1(or683)* reduces the number of embryos with an anterior PAR-2 domain. PLK-1 may phosphorylate and inhibit GSP-2 directly. However, our efforts to mutagenize predicted PLK-1 phosphorylation sites of GSP-2 into non-phosphorylatable ones did not show any phenotype consistent with an upregulation of GSP-2.

**Figure 7:**
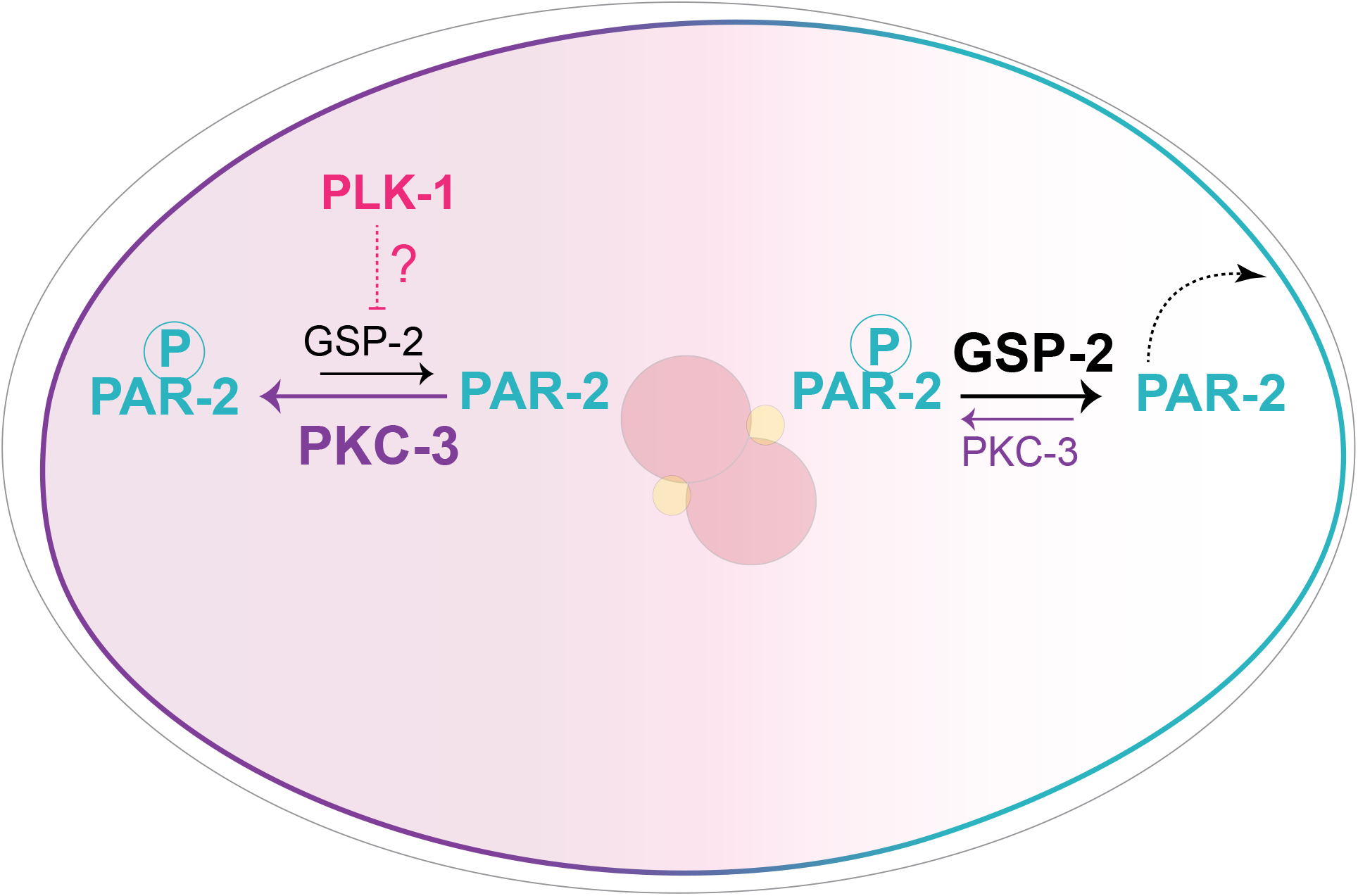
PP1 phosphatases antagonize PKC-3. Schematic representation of a one-cell stage embryo with anterior PAR proteins (in magenta) and posterior PAR proteins (in cyan). At the anterior, phosphorylation of PAR-2 by PKC-3 inhibits PAR-2 cortical localization, whereas at the posterior dephosphorylation of PAR-2 by GSP-2 allows PAR-2 posterior cortical localization. PLK-1 (in pink) downregulates, directly or indirectly, the activity of GSP-2 in the anterior of the embryo.

GSP-1 and GSP-2 are catalytic subunits of the protein phosphatase PP1. Catalytic subunits bind to regulatory subunits and these, in turn, regulate substrate specificity and the activity of the holoenzyme (Aggen et al., 2000). Therefore, we cannot rule out the hypothesis that PLK-1 might act on the yet to be identified regulatory subunits.

Further studies are needed to understand if PLK-1 regulates PP1 activity and whether the regulation is direct or indirect.

All together our data suggest a model in which the proper balance between the activity of phosphatases and PKC-3 is crucial to properly establish polarity in one-cell embryos (Fig. 7).

## Materials and Methods

### Strains

The *C. elegans* strains used in this work are listed in Table S1. Worms were maintained on NGM plates seeded with OP50 bacteria, using standard methods (Brenner, 1974). Thermosensitive strains such as *pkc-3(ne4246)*, *pkc-3(ne4246)*; *gfp::par-2* and *plk-1(or683)* were maintained at 15°C. All the other strains were maintained at 22°C.

Mutant strains were generated using CRISPR/Cas9 technology as described in (Arribere et al., 2014). Single guide RNAs, repair templates to generate the mutant, PCR primers used to detect and sequence the mutation and enzymes used for the screening are listed in Table S2-4. The ZU316 was backcrossed three times. For the ZU297, two independent clones were analyzed.

### RNA interference

Clones from the Ahringer feeding library (Ahringer, 2006; Kamath et al., 2003) were used when available (see Table S5). For the control, in the injection experiments, we used the clone C06A6.2 previously found in the laboratory to have no effect on the early embryonic cell division and to be 100% viable (Bondaz et al., 2019). For GSP-2, a DNA fragment was amplified from cDNA using Gateway-compatible oligonucleotide primers for Gateway-based-cloning into the final pDEST-L4440 vector (forward primer, GGGGACAAGTTTGTACAAAAAAGCAGGCT-GTGACGTGCACGGACAATAC, reverse primer, GGGGACCACTTTGTACAAGAAAGCTGGGT-CTGGTGAGCTCTGCAAATC). To produce dsRNA for injections, the Promega Ribomax RNA production system was used.

dsRNA was injected in L4/young hermaphrodite adults which were incubated at 20°C; embryos from injected hermaphrodites were analyzed after 24-28h (Fig. 5 B) and 18-20h (Figs. 2 C, D, and E, 4 E and F) after injection.

For the depletion of PKC-3 in the *gfp::par-2(RAFA)* mutant strain, which is embryonic lethal, L4/young adult worms were singled on OP50 seeded-NGM plates, let them lay few eggs, and subsequently injected with *pkc-3* dsRNA. Injected worms were transferred to a new OP50 seeded-NGM plate. Homozygous mutant worms were recognized by looking at the un-hatched progeny in the original plates of non-injected worms. For the co-depletion of GSP-1 and GSP-2, a mixture of 1:1 dsRNAs was injected (Figs. 2 C, D and E).

RNA interference by feeding was performed using feeding plates with 1mM IPTG for the *pkc-3(ne4246)* experiments and 3mM IPTG for the *plk-1(or683)* experiments. The empty L4440 vector was used as control. In Figs. 1 B, C and D, 2 B, S1 and S2 (feeding of the *pkc-3(ne4246)* strains and controls) worms were incubated at 24°C since at 25°C most *pkc-3(ne4246)* worms were sterile. L4 larvae were added to RNAi feeding plates and incubated for 24h. For the experiment in Figure 6 (feeding of the *plk-1(or683)* strain and N2) L1 worms were incubated at 15°C until the adult stage.

### Live imaging of embryos

Adult hermaphrodites were dissected on a coverslip into a drop of Egg Buffer (118 mM NaCl, 48 mM KCl, 2 mM CaCl_2_, 2 mM MgCl_2_, and 25 mM Hepes pH 7.5). Embryos were mounted on a 3% agarose pad. Time lapse recordings (frames captured every 10 seconds) were performed using a Nikon ECLIPSE Ni-U microscope, equipped with a Nikon DS-U3 Digital Camera and a 60X objective. For Fig. 1 D it was also used a Leica DM6000 microscope, equipped with a DFC 360 FX camera and a 63X objective. Images of embryos were taken every 10 seconds.

### Immunostaining of embryos and image acquisition

For staining of embryos, 20 gravid hermaphrodites were dissected in a drop of M9 (86 mM NaCl, 42 mM Na2HPO4, 22 mM KH2PO4, and 1 mM MgSO4) on an epoxy slide square (Thermo Fisher Scientific), previously coated with 0.1% poly-L-lysine. A 22X40-mm coverslip was added crosswise on the slide to squash the embryos. The slides were transferred on a metal block on dry ice for at least 10 min. Afterward, the coverslip was removed before fixation for 20 min in methanol. Immunostaining was performed as described in (Spilker et al., 2009). The slides were transferred to a solution of Phosphate Buffered Saline (PBS) plus 0.2% Tween 20 (PBST) and 1% BSA for 20 min. The slides were incubated with primary antibodies diluted in PBST with 1% BSA overnight at 4°C (Rabbit anti PLK-1 (1:500) (Tavernier et al., 2015), Rabbit anti-PAR-2 (1:200) (Labbé et al., 2006), Mouse anti-Tubulin (1:100) (Sigma)).

After two washes of 10 min each in PBST, slides were incubated for 45 min at 37°C with a solution containing secondary antibodies (4 μ/ml Alexa Fluor 488– and/or 568– coupled anti-rabbit or anti-mouse antibodies) and 1 μg/ml DAPI to visualize DNA. Slides were then washed two times for 10 min in PBST before mounting using Mowiol [Calbiochem, 475904], 0.2 M Tris pH 8.5, and 0.1% DABCO).

Images were acquired using a Nikon ECLIPSE Ni-U microscope, equipped with a Nikon DS-U3 Digital Camera, and using a 60X objective.

### Yeast two-hybrid assay

The interaction between PAR-2 and GSP-1/-2 was assessed using a GAL4-based system (Gateway, Invitrogen) using the MAV203 yeast strain. Full-length cDNAs of GSP-1 and GSP-2 were fused to the GAL4 DNA binding domain (Bait plasmid). A PAR-2 (1-335) fragment both wild type and mutant (RAFA) were fused to the GAL4 activation domain (Prey plasmid). The PAR-2 wild type and mutant fragments and the GSP-1 and GSP-2 full length were first cloned into the pDONR201 and subsequently transferred to the pDEST22 vector (GAL4AD) and pDEST32 (GAL4DBD) respectively using Gateway technology. Mutations were inserted by Pfu site directed mutagenesis. A list of plasmids and primers used for the Y2H is provided in Table S6 and S7 respectively. Transformants were selected on Synthetic Defined (SD) medium (lacking Leucine and Tryptophan) plates. The interactions were tested by spotting single colonies containing the desired plasmids on medium lacking Leucine, Tryptophan and Histidine and containing 50 mM of 3AT (3-amino-1,2,3-triazole, Sigma). Picture of the plates were taken using the Fusion FX6 EDGE Imaging System (Vilber) equipped with an Evo-6 Scientific Grade CCD camera.

### Embryonic lethality

To count the embryonic lethality, young adult worms were singled onto NGM plates seeded with OP50 and incubated 24h at 24°C (Figs. 1 B and 2 B) and 20°C (Fig. S3 A and for the *gfp::par-2(L165V)* mutant). After 24 hours, the adult worms were removed and the plates were again incubated at 24°C and 20°C respectively for 24 hours. The ratio between the un-hatched embryos over the total F1 progeny (un-hatched embryos and larvae) was used to calculate the percentage of embryonic lethality.

### Image analysis and measurement

#### PAR-2 cortical intensity line profile (Figs. 2 D and 4 B)

The line profile of cortical PAR-2 was measured in one-cell stage embryos at pronuclear meeting. A segment of 5- pixel-wide and constant length, centered at the posterior cortex of the embryo and positioned with an angle of 90°C to the cortex was traced. For each embryo, the average of three segment (upper, center and lower posterior cortex) was used for quantification. The segment length was normalized to 1 and the line profile of each embryo was normalized to the average of the value in the cytoplasm at position 0 of the segment traced.

#### Cortical PAR-2 size measurement (Figs. 2 A, 2 E, S2 B, S3 C)

The size of cortical *gfp::par-2* domain was determined by measuring the length of the PAR-2 domain normalized for the total perimeter of the embryo at pronuclear meeting. The perimeter of the embryo and the length of the PAR-2 domain were traced manually by using ImageJ software. The length of the PAR-2 domain is represented as a percentage of the total perimeter of the embryo.

#### Cortical PAR-2 intensity (Figs. 4 F and S3 B)

The mean intensity of cortical *gfp::par-2* was determined by tracing a line of 5-pixel-wide along the posterior cortex at pronuclear meeting. The mean intensity of the posterior cytoplasm was obtained making a square of fixed area. For Fig. 4 F the ratio between the PAR-2 intensity at the cortex over the one in the cytoplasm is plotted, whereas for Fig. S3 B the mean intensity of the cytoplasm was subtracted from the mean intensity of cortical PAR-2.

#### Cell cycle timing measurement (Figs. 1 D and 4 D)

Cell cycle length was measured from the onset of Nuclear Envelope Breakdown (NEBD) in P0 to the onset of NEBD in AB and P1. NEBD was measured at the time of nuclear membrane disappearance.

#### AB/P1 ratio measurement (Fig. S1 A)

AB and P1 length were measure at the time of cytokinesis furrow’s ingression and then the ratio AB/P1 was calculated. Value equal to 1 indicates symmetry, whereas value >1 indicates wild type asymmetry with AB bigger than P1.

#### PLK-1 asymmetry measurement (Fig. S1 B)

The area of the AB and P1 cells was determined manually by using ImageJ software and the intensity of PLK-1 was measured. For the nucleus intensity, a circle around the nucleus was drawn. The intensity of the nucleus was subtracted from the intensity of PLK-1 of each cell. PLK-1 ratio was determined by dividing. PLK-1 ratio was determined by dividing the mean of PLK-1 intensity in AB over the mean of PLK-1 intensity in P1. Value equal to 1 indicates symmetric localization of PLK-1 in AB and P1, whereas value >1 indicates wild type asymmetry with PLK-1 more enriched in AB compared to P1.

#### Cleavage furrow position measurement (Fig. S3 D)

The total length of the embryo and the length from the anterior pole to the cleavage furrow were measured manually using ImageJ software. The cleavage furrow position was determined as ratio of the length from the anterior pole to the cleavage furrow over the total length of the embryo.

### Protein phosphatase motif identification

The PP1 docking motif was identified using the Eukaryotic Linear Motif resource for Functional Sites in Proteins (http://elm.eu.org).

### Statistical analysis

Statistical analysis was performed using GraphPad Prism 9. Details regarding the statistical test, the sample size, the experiment number and the meaning of error bars are provided for each experiment in the corresponding figure legend, in the results and/or in the method details. Refer also to Table S8-9.

Significance was defined as: ns, p > 0.05, *p < 0.05, **p < 0.01, ***p < 0.001, ****p < 0.0001.

### Online supplemental material

Fig. S1 describes cell size asymmetry and PLK-1 localization in *pkc-3* temperature sensitive mutant and in GSP-2 depleted embryos. Fig. S2 describes the phenotype of *gsp-1(RNAi)* embryos. Fig. S3 compares the phenotypes (lethality, PAR-2 intensity and domain size and cell size) of *gfp::par-2* and *gfp::par-2(RAFA)* embryos. Video 1 and 3 show the first division and *gfp::par-2* localization of *pkc-3(ne4246)* and *pkc-3(ne4246); gsp-2(RNAi)* (Video 1) and *pkc-3(ne4246); gsp-1(RNAi)* (Video 3) embryos. Video 2 shows the first and second division of *pkc-3(ne4246)* and *pkc-3(ne4246); gsp-2(RNAi)* embryos. Video 4 shows the first division and *gfp::par-2* localization of *gsp-1/-2(RNAi)* and control embryos. Video 5 shows the PAR-2 localization in *gfp::par-2(RAFA)* embryos and control. Video 6 shows the first and second division of *gfp::par-2(RAFA)* embryos and control. Video 7 shows the PAR-2 localization in *gfp::par-2(L165V)* embryos and control. Table S1 shows the genotypes of strains used in this study. Tables S2-S4 show all the reagents used for CRISPR. Table S5 summarizes the clones used for RNA interference. Tables S6 and S7 summarize the plasmids used for the two-hybrid experiment. Tables S8 and S9 summarize the statistical analyses used in this study.

## Acknowledgments

We would like to thank N. Goehring (Francis Crick Institute) for reagents and discussions. We thank present and past members of the Gotta laboratory for discussions, suggestions and feedback on the manuscript. A special thank to Sofia Barbieri for help with the statistical analysis. We thank Patrick Meraldi, Florian Steiner and their laboratories for fruitful discussions and comments on the manuscript. Some strains were provided by the CGC, which is funded by the NIH office of research infrastructure program (P40OD010440). Work in the laboratory of MG is funded by the SNF (grant number 31003A_175850) and by the University of Geneva.

The authors declare no competing financial interests.

## Authors contribution

I. Calvi and M. Gotta conceived the project. I. Calvi performed all experiments, with the exception of the experiment in Fig. 3 B, which was performed by F. Schwager. I. Calvi and M. Gotta analyzed and interpreted all the results and wrote the manuscript.

