## Supplementary Information for "PP1 phosphatases control PAR-2 localization and polarity establishment in *C. elegans* embryos"

**Figure S1**

**A**

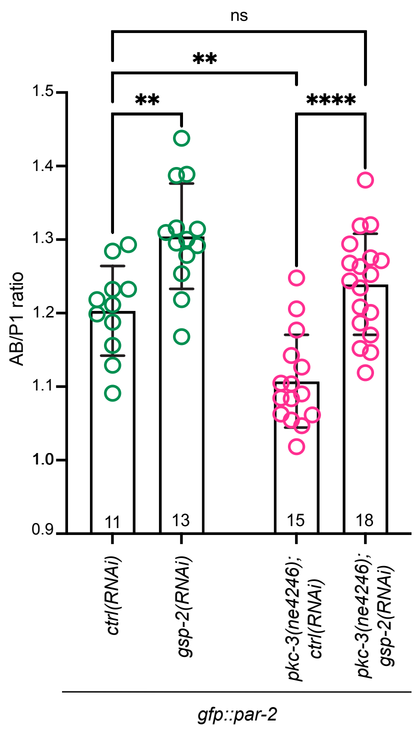

ns
**
**
****
1.5
1.4
1.3
1.2
1.1
1.0
0.9
AB/P1 ratio
11
13
15
18
ctrl(RNAi)
gsp-2(RNAi)
pkc-3(ne4246); ctrl(RNAi)
pkc-3(ne4246); gsp-2(RNAi)
gfp::par-2


**B**

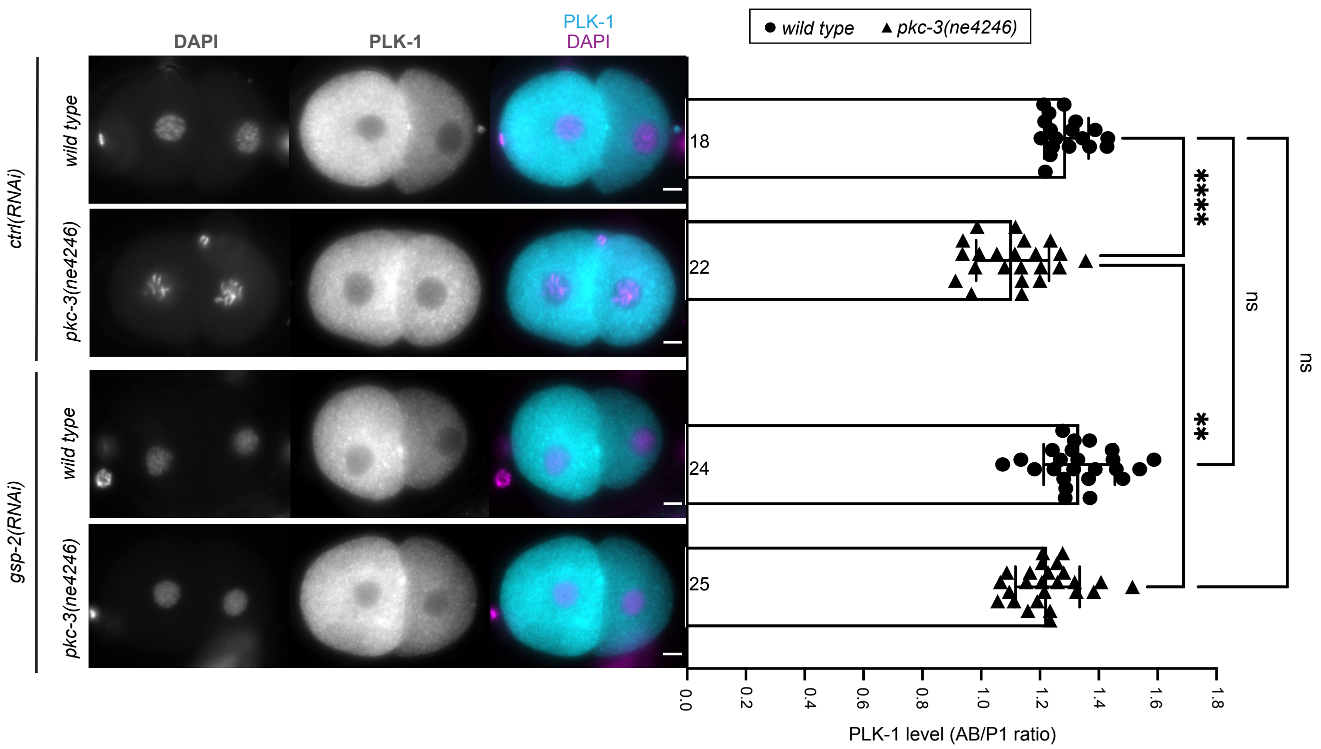

DAPI
PLK-1
PLK-1
DAPI
wild type
pkc-3(ne4246)
ctrl(RNAi)
gsp-2(RNAi)
● wild type ▲ pkc-3(ne4246)
18
22
24
25
****
**
ns
ns
0.0
0.2
0.4
0.6
0.8
1.0
1.2
1.4
1.6
1.8
PLK-1 level (AB/P1 ratio)

**Figure S2**

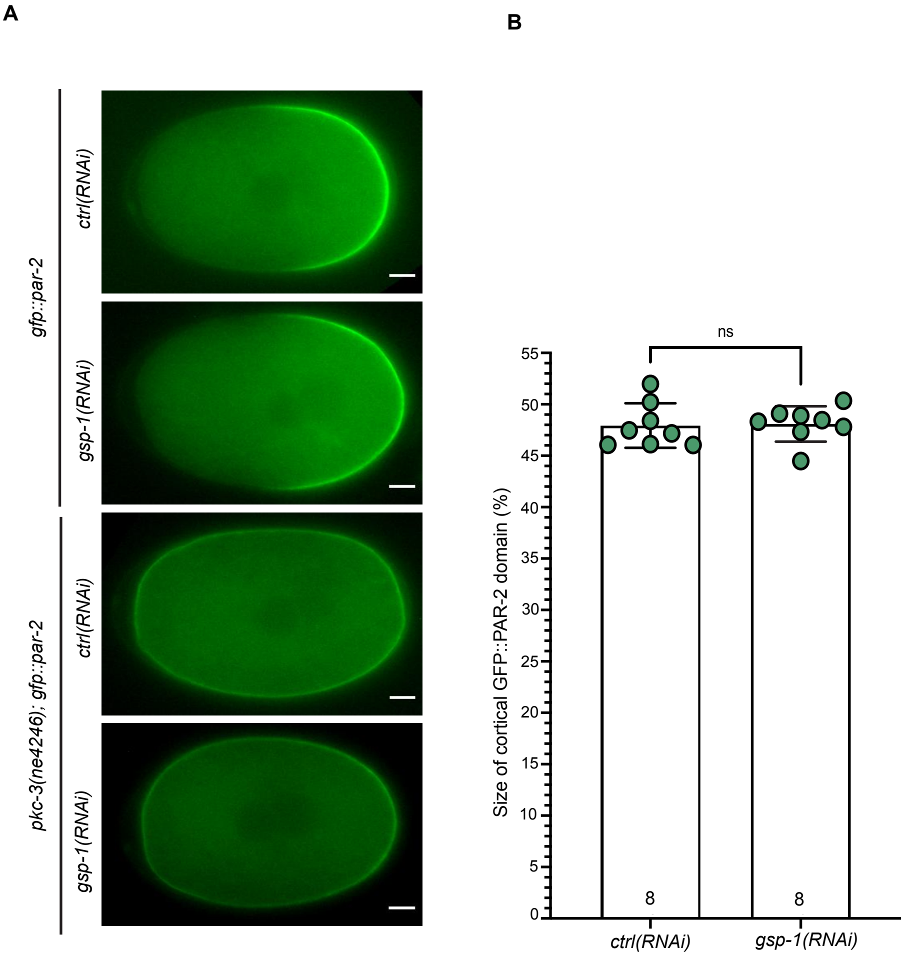

A
B
gfp::par-2
pkc-3(ne4246); gfp::par-2
ctrl(RNAi)
gsp-1(RNAi)
ctrl(RNAi)
gsp-1(RNAi)
ns
Size of cortical GFP::PAR-2 domain (%)
0
5
10
15
20
25
30
35
40
45
50
55
8
8
ctrl(RNAi)
gsp-1(RNAi)

**Figure S3**

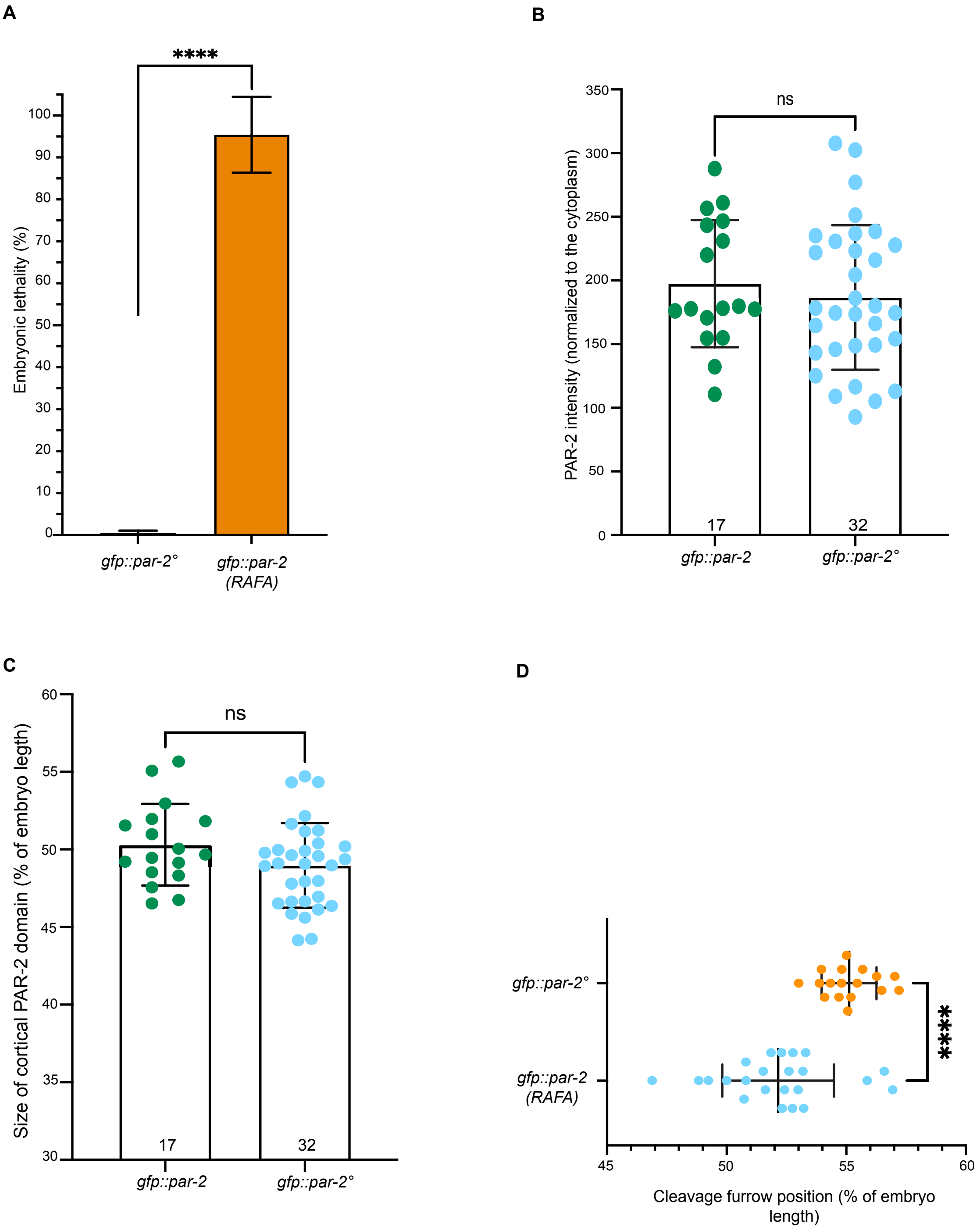
A
****
Embryonic lethality (%)
0
10
20
30
40
50
60
70
80
90
100
gfp::par-2°
gfp::par-2
(RAFA)
B
ns
PAR-2 intensity (normalized to the cytoplasm)
0
50
100
150
200
250
300
350
17
32
gfp::par-2
gfp::par-2°
C
ns
Size of cortical PAR-2 domain (% of embryo legth)
30
35
40
45
50
55
60
17
32
gfp::par-2
gfp::par-2°
D
gfp::par-2°
gfp::par-2
(RAFA)
****
45
50
55
60
Cleavage furrow position (% of embryo
length)

**SUPPLEMENTARY INFORMATION**

**Supplementary figures legends**

**Figure S1: *gsp-2(RNAi)* rescues AB/P1 asymmetry and PLK-1 localization in the pkc-3(ne4246) mutant allele**

(**A)** Quantification of cell size asymmetry (AB/P1 ratio). AB/P1 ratio equal to 1 correspond to cell symmetry, values higher or lower than 1 correspond to cell asymmetry. Numbers inside the bars indicate sample size. N=4. Mean is shown and error bars indicate SD. The P-values were determined using 2way ANOVA "Tukey's multiple comparisons test".

**(B)** Representative two-cell stage images of fixed embryos of the indicated genotypes. DNA (DAPI) is in magenta and PLK-1 in cyan. Right: quantification of the PLK-1 levels (AB/P1 ratio) in the two-cell stage fixed embryos. Numbers inside the bars indicate sample size. N=2. Mean is shown and error bars indicate SD. The P-values were determined using 2way ANOVA "Tukey's multiple comparisons test". ns p > 0.05, **p < 0.01, ****p < 0.0001. Scale bars, 5 µm. Anterior is to the left and posterior to the right.

**Figure S2: *gsp-1(RNAi)* does not rescue PAR-2 localization in the *pkc-3(ne4246); gfp::par-2* strain and does not affect PAR-2 size domain in the *gfp::par-2* strain**

**(A)** Representative midsection frames of time-lapse videos of *gfp::par-2* and *pkc-3(ne4246); gfp::par-2* one-cell embryos at pronuclear meeting, comparing *ctrl(RNAi)* and *gsp-1(RNAi).* In *gfp::par-2*; *ctrl(RNAi)* and *gfp::par-2*; *gsp-1(RNAi),* PAR-2 localizes at the posterior cortex (n=8 and n=8 respectively). In *pkc-3(ne4246); gfp::par-2*, *ctrl(RNAi)* and *gsp-1(RNAi)* PAR-2 signal is detected uniformly at the cortex (n=10 and n=13 respectively). N=3. **(B)** Quantification of the *gfp::par-2* size domain in live zygotes at pronuclear meeting, comparing *ctrl(RNAi)* and *gsp-1(RNAi).* Numbers inside the bars indicate sample size. N=3. The P-values were determined using unpaired Student's t-test. Mean is shown and error bars indicate SD. ns p > 0.05. Scale bars, 5 μm. Anterior is to the left and posterior to the right.

**Figure S3: The PP1 motif in PAR-2 is crucial for proper cell polarity**

**(A)** Embryonic lethality of *gfp::par-2°* (refer also later to a homozygous worms expressing wild-type *gfp::par-2* and heterozygous worms expressing wild type *gfp::par-2* from one allele and mutant *gfp::par-2(RAFA)* from the other allele) and *gfp::par-2(RAFA)* homozygous mutant strain. The reported values correspond to the percentage of un-hatched embryos over the total progeny (larvae and un-hatched embryo). *gfp::par-2°*, n=2469, *gfp::par-2(RAFA)*, n=1028. N=4. Mean is shown and error bars indicate SEM. The P-values were determined using Student's t-test. **(B)** Measurement of the PAR-2 cortical intensity in the wild type *gfp::par-2* and in the *gfp::par-2°* embryos at pronuclear meeting. Numbers inside the bars indicate sample size. N=2. **(C)** Quantification of the PAR-2 size domain in live zygotes at pronuclear meeting, comparing *gfp::par-2* and *gfp::par-2°.* Numbers inside the bars indicate sample size. N=2. Mean is shown and error bars indicate SD. The P-values were determined using unpaired Student's t-test. **(D)** Quantification of the cleavage furrow position in *gfp::par-2°*and *gfp::par-2(RAFA)*. *gfp::par-2(RAFA)* mutant show cleavage furrow position more towards the anterior compared with *gfp::par-2°* (n=23 and n=17 respectively). N=6. Mean is shown and error bars indicate SD. The P-values were determined using unpaired Student's t-test. ns p > 0.05, ****p < 0.0001. Scale bars, 5 μm. Anterior is to the left and posterior to the right.

### Videos legends

**Video 1: *gsp-2(RNAi)* rescues PAR-2 posterior cortical localization in the *pkc-3(ne4246); gfp ::par-2* embryos**

*gfp::par-2* and *pkc-3(ne4246); gfp::par-2* embryos treated either with *ctrl(RNAi)* or *gsp-2(RNAi)*. Acquisition of midplane fluorescent images begins during the early establishment phase, and frames are captured every 10 s. In *gfp::par-2* embryos (both *ctrl(RNAi)* and *gsp-2(RNAi)*) PAR-2 localizes at the posterior cortex in the one-cell stage embryo and in P1 in the two-cell stage embryo (n=11 and n=13 respectively). In *pkc-3(ne4246); gfp::par-2*, *ctrl(RNAi)* PAR-2 is uniformly distributed at the cortex in both one and two cell stage embryo (n=15). After depletion of GSP-2, *pkc-3(ne4246); gfp::par-2* embryos showed PAR-2 posterior localization in one-cell stage embryo and in the P1 cell (n=18). N=4. Anterior is to the left and posterior to the right. Scale bar: 5 µm.

**Video 2: Depletion of GSP-2 restores the cell cycle asynchrony between AB and P1 in *pkc-3(ne4246)* embryos**

*Control* and *pkc-3(ne4246)* embryos after *ctrl(RNAi)* and *gsp-2(RNAi)*. Acquisition of midplane DIC images from NEBD in P0 to four-cell stage. Frames are captured every 10s. Depletion of *gsp-2(RNAi)* in *pkc-3(ne4246)* embryos restores AB and P1 asynchrony. These four examples were chosen because they had cell cycle time close to the mean of each population. *ctrl(RNAi)* n=12, *gsp-2(RNAi)* n=10, *pkc-3(ne4246); ctrl(RNAi)* n=10 and *pkc-3(ne4246); gsp-2(RNAi)* n=18. N=3. Anterior is to the left and posterior to the right. Scale bar: 5 µm.

**Video 3**: ***gsp-1(RNAi)* does not rescue PAR-2 posterior cortical localization in the *pkc-3(ne4246); gfp ::par-2* embryos**

*gfp::par-2* and *pkc-3(ne4246); gfp::par-2* embryos treated either with *ctrl(RNAi)* or *gsp-1(RNAi)*. Acquisition of midplane fluorescent images begins during the early establishment phase, and frames are captured every 10 s. In *gfp::par-2* embryos (both *ctrl(RNAi)* and *gsp-2(RNAi) n=8*) PAR-2 localizes at the posterior cortex in the one-cell stage embryo and in P1 in the two-cell stage embryo. In *pkc-3(ne4246); gfp::par-2*, *ctrl(RNAi)* and *gsp-1(RNAi)* PAR-2 is uniformly distributed at the cortex in both one and two cell stage embryo (n=10 and n=13 respectively). N=3. Anterior is to the left and posterior to the right. Scale bar: 5 µm.

**Video 4: GSP-1 and GSP-2 co-depletion impairs PAR-2 posterior cortical localization**

*gfp::par-2* embryos treated with *ctrl(RNAi)*, n=10, *ctrl(RNAi); gsp-1(RNAi)*, =9, *ctrl(RNAi); gsp-2(RNAi),* n=11*,* and *gsp-1/-2(RNAi) class I*, n=11, and *class II,* n=3. N=3. Acquisition of midplane fluorescent images begins during the early establishment phase, and frames are captured every 10 s. In *gfp::par-2; ctrl(RNAi)*, *ctrl(RNAi); gsp-1(RNAi)*, *ctrl(RNAi); gsp-2(RNAi)* embryos, PAR-2 localizes at the posterior cortex in the one-cell stage embryo and in P1 in the two-cell stage embryo. In *gfp::par-2 gsp-1/-2(RNAi) class I* embryos PAR-2 is detected in the cytoplasm, whereas in *gsp-1/-2(RNAi) class II* embryos PAR-2 is weakly detected at the cortex compared to the *ctrl(RNAi).* N=3. Anterior is to the left and posterior to the right. Scale bar: 5 µm.

**Video 5: The PP1 motif in PAR-2 is required for its localization at the posterior cortex**

*gfp::par-2°* (the symbol ° indicates a mixture between wild-type *gfp::par-2* homozygous worms expressing wild-type *gfp::par-2* and heterozygous worms expressing wild type *gfp::par-2* from one allele and mutant *gfp::par-2(RAFA)* from the other allele) and *gfp::par-2(RAFA)* homozygote mutant embryos. Acquisition of midplane fluorescent images begins during the early establishment phase, and frames are captured every 10s. In *gfp::par-2°* embryos PAR-2 localizes at the posterior cortex in the one-cell stage embryo and in P1 in the two-cell stage embryo (n=16). In *gfp::par-2(RAFA)* embryos, PAR-2 is mostly detected in the cytoplasm (n=17). N=6. Anterior is to the left and posterior to the right. Scale bar: 5 μm.

**Video 6: Two-cell stage *gfp::par-2(RAFA)* embryos divide synchronously**

Acquisition of midplane DIC images from NEBD in P0 to four-cell stage for *gfp::par-2°* and *gfp::par-2(RAFA)* (n=14 and n=18 respectively). Frames are captured every 10s. In *gfp::par-2(RAFA)* embryos, AB and P1 divide synchronously. These four examples were chosen because they had cell cycle time similar to the mean of each population. N=6. Anterior is to the left and posterior to the right. Scale bar: 5 μm.

**Video 7: *gfp::par-2(L165V)* mutant embryos show aberrant PAR-2 cortical localization**

*gfp::par-2* and *gfp::par-2(L165V)* embryos. Acquisition of midplane fluorescent images begins during the early stage, and frames are captured every 10s. In *gfp::par-2(L165V)* embryos PAR-2 localizes at both the anterior and posterior cortex in the early phase and is removed from the anterior in most of the embryos during the first cell division compared to *gfp::par-2* where PAR-2 show a normal localization at the posterior cortex (n=25 and n=14 respectively). N=4. Anterior is to the left and posterior to the right. Scale bar: 5 μm.

**Table S1**

| **Strain name** | **Genotype** | **Source/Reference** | **Description** |
|---|---|---|---|
| **N2** | *Wild type* | *Caenorhabditis* Genetics Center (*CGC*) | |
| **EU1441** | *plk-1(or683) III* | (O'Rourke et al., 2011) | |
| **KK1273** | *par-2(it328[gfp::par-2]) III* | *Caenorhabditis* Genetics Center (*CGC*) | |
| **NWG0124** | *pkc-3(ne4246) II*; par-2 *(it328[gfp::par-2]) III* | Courtesy of Nathan Goehring | |
| **WM150** | *pkc-3*(*ne4246*) *II* | (Fievet et al., 2013) | |
| **ZU297** | *par-2(it328[gfp::par-2]) (L165A, F167F) III* | This study | Strain generated by CRISPR/Cas9 inserting two point mutations in aa position 165 and 167. The background strain is KK1273. |
| **ZU316** | *par-2(it328[gfp::par-2]) (L165V) III* | This study | Strain generated by CRISPR/Cas9 inserting one point mutation in aa position 165. The background strain is KK1273. |

### Table S2

| Strain | sgRNA sequences (5'→3') |
|---|---|
| ZU297 | 1. F : TCTTG TGACGAGAAGCCTGTCACAA<br>R : AAAC TTGTGACAGGCTTCTCGTCA C<br><br>2. F : TCTTG ACGCTTATTCTTCTCAATTG<br>R : AAAC CAATTGAGAAGAATAAGCGT C |
| ZU316 | Same as ZU297 |

In the sgRNA sequences, underlined nucleotides define the overhang complementary to the recipient guide RNA vector (pRB1017).

### Table S3

| Strain | Repair template sequence (5'→3') |
|---|---|
| ZU297 | GGAAATTCCGGCTCAAAAAATGACGAGAAGCCTGTCACAA**CGc**TCTAAACGACGC*gc*ATTC*gc*CTCAATTG**CcG**GCCCAACGATTTGCTCGAGgtagatggctggaaaagagctg |
| ZU316 | GGAAATTCCGGCTCAAAAAATGACGAGAAGCtTGTCACAA**CGc**TCTAAACGACGC*g*TATTCTTCTCAATTG**CcG**GCCCAACGATTTGCTCGAGgtagatggctggaaaagagctg |

The underlined sequence is targeted by the sgRNA.

For ZU297 strain: In bold the **PAM** sites silently mutated (originally CGG mutated in **CGc** and CGG mutated in **CcG)**. In italic the *nucleotide substitution* (TTA in gcA and TTC in gcC).

For ZU316 strain: In bold the **PAM** sites silently mutated (originally CGG mutated in **CGc** and CGG mutated in **CcG**). In italic the *nucleotide substitution* (TTA in gTA).

### Table S4

| Strain | PCR primers (5'→3') | Restriction enzyme used and PCR product size |
|---|---|---|
| ZU297 | F : GGTCCGAAAATTGAAAAACC<br>R : CGAAAATTAGCATTTTTCGGG | BsmI enzyme (NEB) was used for the screening. The sequence recognized by the enzyme is in the 2nd sgRNA sequence (3'..CTTACGN..5').<br>The PCR product is ~ 1200bp. After BsmI digestion, fragments of ~850bp and ~350bp are visible in the homozygote mutant. |
| ZU316 | Same as ZU297 | The HindIII enzyme (NEB) was used for the screening. The sequence recognized by the enzyme is in the 1st sgRNA sequence (3'..AAGCTT..5').<br>The PCR product is ~ 1200bp. After HindIII digestion, fragments of ~865bp, ~180bp and ~130bp are visible in the homozygote mutant. |

### Table S5

| Target gene | Clone names from Ahringer library | Oligonucleotides sequence (5'→3') | |
|---|---|---|---|
| *control* | C06A6.2 | T7 primers | CGTAATACGACTCACTATAG |
| *gsp-1* | F29F11.6 | T7 primers | CGTAATACGACTCACTATAG |
| *gsp-2* | | Forward<br>GTGACGTGCACGGACAATAC | Reverse<br>CTGGTGAGCTCTGCAAATC |
| *pkc-3* | F09E5.1 | T7 primers | CGTAATACGACTCACTATAG |

### Table S6

| Plasmid | Backbone | Insert | Tag | Reference |
|---|---|---|---|---|
| - | pDONR201 | Empty | - | Invitrogen |
| pMG1262 | pDONR201 | PAR-2b cDNA (1-335aa) | - | This study |
| pMG126 | pDONR201 | PAR-2b cDNA (1-335aa, L165A, F167A)) | - | This study |
| pMG1259 | pDONR201 | GSP-2 cDNA (f.l.) | - | This study |
| pMG1463 | pDONR201 | GSP-1 cDNA (f.l.) | - | This study |
| - | pDEST AD | Empty | Y2H AD | Invitrogen |
| pMG726 | pDEST AD | PAR-2b cDNA (1-335aa) | Y2H AD | This study |
| pMG430 | pDEST AD | PAR-2b cDNA (1-335aa, L165A, F167A) | Y2H AD | This study |
| - | pDEST DBD | Empty | Y2H DBD | Invitrogen |
| pMG1264 | pDEST DBD | GSP-2 cDNA (f.l.) | Y2H DBD | This study |
| pMG1464 | pDEST DBD | GSP-1 cDNA (f.l.) | Y2H DBD | This study |

### Table S7

| Plasmid | Primers (5'→3') |
|---|---|
| pMG1262 | F: GGGGACAAGTTTGTACAAAAAAGCAGGCT*TG*ATGACGGATTTAGACACTTCATC<br>R: GGGGACCACTTTGTACAAGAAAGCTGGGT*G*CTAGCTGGATTTGGTCGGCTTCT |
| pMG126 | F: CGACGCgcATTCgcCTCAATTG<br>R: CAATTGAGgcGAATgcGCGTCG |
| pMG1259 | F: GGGGACAAGTTTGTACAAAAAAGCAGGCT*TG*ATGGACGTAGAAAAGCTTAATC<br>R: GGGGACCACTTTGTACAAGAAAGCTGGGT*G*TTACTTCTTGGCACCCTTCTTT |
| pMG1463 | F: GGGGACAAGTTTGTACAAAAAAGCAGGCT*TG*ATGTCGAACGATGGAGATTTAAAC<br>R: GGGGACCACTTTGTACAAGAAAGCTGGGT*G*TTACTTCTTACCAGCAGTCGTTC |
| pMG1487 | F: CTAAACGACGCgTATTCTTCTC<br>R: GAGAAGAATAcGCGTCGTTTAG |

Underlined sequences are the Gateway overhang.

In italic the *nucleotides* added to keep the ORF.

In lower case the nucleotides mutation.

Fievet, B. T., Rodriguez, J., Naganathan, S., Lee, C., Zeiser, E., Ishidate, T., Shirayama, M., Grill, S., & Ahringer, J. (2013). Systematic genetic interaction screens uncover cell polarity regulators and functional redundancy. *Nat Cell Biol*, *15*(1), 103-112. https://doi.org/10.1038/ncb2639

O'Rourke, S. M., Carter, C., Carter, L., Christensen, S. N., Jones, M. P., Nash, B., Price, M. H., Turnbull, D. W., Garner, A. R., Hamill, D. R., Osterberg, V. R., Lyczak, R., Madison, E. E., Nguyen, M. H., Sandberg, N. A., Sedghi, N., Willis, J. H., Yochem, J., Johnson, E. A., & Bowerman, B. (2011). A survey of new temperature-sensitive, embryonic-lethal mutations in C. elegans: 24 alleles of thirteen genes. *PLoS One*, *6*(3), e16644. https://doi.org/10.1371/journal.pone.0016644

### Table S8

| Figure | Genotypes compared | Summary | p-value | Test |
|---|---|---|---|---|
| 1B | *ctrl(RNAi)*:*gfp::par-2* vs. *ctrl(RNAi)*:*pkc-3(ne4246); gfp::par-2* | **** | <0.0001 | 2way-anova "Tukey's multiple comparisons test" |
| | *ctrl(RNAi)*:*gfp::par-2* vs. *gsp-2(RNAi)*:*gfp::par-2* | ns | 0.9971 | 2way-anova "Tukey's multiple comparisons test" |
| | *ctrl(RNAi)*:*gfp::par-2* vs. *gsp-2(RNAi)*:*pkc-3(ne4246); gfp::par-2* | * | 0.0283 | 2way-anova "Tukey's multiple comparisons test" |
| | *ctrl(RNAi)*:*pkc-3(ne4246); gfp::par-2* vs. *gsp-2(RNAi)*:*gfp::par-2* | **** | <0.0001 | 2way-anova "Tukey's multiple comparisons test" |
| | *ctrl(RNAi)*:*pkc-3(ne4246); gfp::par-2* vs. *gsp-2(RNAi)*:*pkc-3(ne4246); gfp::par-2* | **** | <0.0001 | 2way-anova "Tukey's multiple comparisons test" |
| | *gsp-2(RNAi)*:*gfp::par-2* vs. *gsp-2(RNAi)*:*pkc-3(ne4246); gfp::par-2* | * | 0.0399 | 2way-anova "Tukey's multiple comparisons test" |
| 1D | *ctrl(RNAi):* AB vs. *ctrl(RNAi)* P1 | **** | <0.0001 | Unpaired Student's t-test |
| | *gsp-2(RNAi):* AB vs. *gsp-2(RNAi):* P1 | **** | <0.0001 | Unpaired Student's t-test |
| | *ctrl(RNAi): pkc-3(ne4246)* AB vs. *ctrl(RNAi): pkc-3(ne4246)* P1 | ns | 0.3406 | Unpaired Student's t-test |

| | | | | |
|---|---|---|---|---|
| | *gsp-2(RNAi): pkc-3(ne4246)* AB vs. *gsp-2(RNAi): pkc-3(ne4246)* P1 | **** | <0.0001 | Unpaired Student's t-test |
| | *ctrl(RNAi): pkc-3(ne4246)* P1 vs. *gsp-2(RNAi): pkc-3(ne4246)* P1 | *** | 0.0006 | Unpaired Student's t-test |
| 2A | *ctrl(RNAi)*:GFP::PAR-2 vs. *gsp-2(RNAi)*:GFP::PAR-2 | **** | <0.0001 | Unpaired Student's t-test |
| 2B | *ctrl(RNAi)*:*gfp::par-2* vs. *ctrl(RNAi)*:*pkc-3(ne4246); gfp::par-2* | **** | <0.0001 | 2way-anova "Tukey's multiple comparisons test" |
| | *ctrl(RNAi)*:*gfp::par-2* vs. *gsp-1(RNAi)*:*gfp::par-2* | ns | 0.2163 | 2way-anova "Tukey's multiple comparisons test" |
| | *ctrl(RNAi)*:*gfp::par-2* vs. *gsp-1(RNAi)*:*pkc-3(ne4246); gfp::par-2* | **** | <0.0001 | 2way-anova "Tukey's multiple comparisons test" |
| | *ctrl(RNAi)*:*pkc-3(ne4246); gfp::par-2* vs. *gsp-1(RNAi)*:*gfp::par-2* | **** | <0.0001 | 2way-anova "Tukey's multiple comparisons test" |
| | *ctrl(RNAi)*:*pkc-3(ne4246); gfp::par-2* vs. *gsp-1(RNAi)*:*pkc-3(ne4246); gfp::par-2* | ns | 0.1584 | 2way-anova "Tukey's multiple comparisons test" |
| | *gsp-1(RNAi)*:*gfp::par-2* vs. *gsp-1(RNAi)*:*pkc-3(ne4246); gfp::par-2* | **** | <0.0001 | 2way-anova "Tukey's multiple comparisons test" |
| 2D | *ctrl(RNAi)*:*gfp::par-2* vs. *ctrl(RNAi); gsp-1(RNAi)*:*gfp::par-2* | ns | 0.9011 | Unpaired Student's t-test |
| | *ctrl(RNAi)*:*gfp::par-2* vs. *ctrl(RNAi); gsp-2(RNAi)*:*gfp::par-2* | *** | 0.0003 | Unpaired Student's t-test |

| | | | | |
|---|---|---|---|---|
| | *ctrl(RNAi)*:*gfp::par-2* vs. g*sp-1/-2(RNAi)*:*gfp::par-2 (class I)* | **** | <0.0001 | Unpaired Student's t-test |
| | *ctrl(RNAi)*:*gfp::par-2* vs. g*sp-1/-2(RNAi)*:*gfp::par-2 (class II)* | **** | <0.0001 | Unpaired Student's t-test |
| 4D | *gfp::par-2°* AB vs. *gfp::par-2°* P1 | ** | 0.0067 | Unpaired Student's t-test |
| | *gfp::par-2(RAFA)* AB vs. *gfp::par-2(RAFA)* P1 | ns | 0.8827 | Unpaired Student's t-test |
| 4F | *ctrl(RNAi)*:*gfp::par-2°* vs. *ctrl(RNAi)*:*gfp::par-2(RAFA)* | **** | <0.0001 | 2way-anova "Tukey's multiple comparisons test" |
| | *ctrl(RNAi)*:*gfp::par-2°* vs. *pkc-3(RNAi)*:*gfp::par-2°* | **** | <0.0001 | 2way-anova "Tukey's multiple comparisons test" |
| | *ctrl(RNAi)*:*gfp::par-2°* vs. *pkc-3(RNAi)*:*gfp::par-2(RAFA)* | **** | <0.0001 | 2way-anova "Tukey's multiple comparisons test" |
| | *ctrl(RNAi)*:*gfp::par-2(RAFA)* vs. *pkc-3(RNAi):gfp::par-2°* | **** | <0.0001 | 2way-anova "Tukey's multiple comparisons test" |
| | *ctrl(RNAi)*:*gfp::par-2(RAFA)* vs. *pkc-3(RNAi):gfp::par-2(RAFA)* | **** | <0.0001 | 2way-anova "Tukey's multiple comparisons test" |
| | *pkc-3(RNAi):gfp::par-2°* vs. *pkc-3(RNAi):gfp::par-2(RAFA)* | ns | 0.0682 | 2way-anova "Tukey's multiple comparisons test" |

### Table S9

| Figure | Genotypes compared | Summary | p-value | Test |
|---|---|---|---|---|
| S1A | *ctrl(RNAi)*:*gfp::par-2* vs. *ctrl(RNAi)*:*pkc-3(ne4246); gfp::par-2* | ** | 0.0017 | 2way-anova "Tukey's multiple comparisons test" |
| | *ctrl(RNAi)*:*gfp::par-2* vs. *gsp-2(RNAi)*:GFP::PAR-2 | ** | 0.0046 | 2way-anova "Tukey's multiple comparisons test" |
| | *ctrl(RNAi)*:*gfp::par-2* vs. *gsp-2(RNAi)*:*pkc-3(ne4246); gfp::par-2* | ns | 0.4117 | 2way-anova "Tukey's multiple comparisons test" |
| | *ctrl(RNAi)*:*pkc-3(ne4246); gfp::par-2* vs. *gsp-2(RNAi)*:*gfp::par-2* | **** | <0.0001 | 2way-anova "Tukey's multiple comparisons test" |
| | *ctrl(RNAi)*:*pkc-3(ne4246); gfp::par-2* vs. *gsp-2(RNAi)*:*pkc-3(ne4246); gfp::par-2* | **** | <0.0001 | 2way-anova "Tukey's multiple comparisons test" |
| | *gsp-2(RNAi)*:*gfp::par-2* vs. *gsp-2(RNAi)*:*pkc-3(ne4246); gfp::par-2* | ns | 0.1017 | 2way-anova "Tukey's multiple comparisons test" |
| S1B | *ctrl(RNAi)*:*wild type* vs. *ctrl(RNAi)*:*pkc-3(ne4246)* | **** | <0.0001 | 2way-anova "Tukey's multiple comparisons test" |
| | *ctrl(RNAi)*:*wild type* vs. *gsp-2(RNAi)*:*wild type* | ns | 0.5913 | 2way-anova "Tukey's multiple comparisons test" |
| | *ctrl(RNAi):wild type* vs. *gsp-2(RNAi)*:*pkc-3(ne4246)* | ns | 0.2454 | 2way-anova "Tukey's multiple comparisons test" |
| | *ctrl(RNAi)*:*pkc-3(ne4246)* vs. *gsp-2(RNAi)*:*wild type* | **** | <0.0001 | 2way-anova "Tukey's multiple comparisons test" |
| | *ctrl(RNAi)*:*pkc-3(ne4246)* vs. *gsp-2(RNAi)*:*pkc-3(ne4246)* | ** | 0.0024 | 2way-anova "Tukey's multiple comparisons test" |
| | *gsp-2(RNAi)*:*wild type* vs. *gsp-2 (RNAi)*:*pkc-3(ne4246)* | ** | 0.0055 | 2way-anova "Tukey's multiple comparisons test" |

| | | | | |
|---|---|---|---|---|
| S2B | *ctrl(RNAi):gfp::par-2* vs. *gsp-1(RNAi):gfp::par-2* | ns | 0.8708 | Unpaired Student's t-test |
| S3A | *gfp::par-2°* vs. *gfp::par-2(RAFA)* | **** | <0.0001 | Unpaired Student's t-test |
| S3B | *gfp::par-2* vs. *gfp::par-2°* | ns | 0.5088 | Unpaired Student's t-test |
| S3C | gfp::par-2 vs. gfp::par-2° | ns | 0.1071 | Unpaired Student's t-test |
| S3D | *gfp::par-2°* vs. *gfp::par-2(RAFA)* | **** | <0.0001 | Unpaired Student's t-test |
